# Genetic mapping of resistance: A QTL and associated polymorphism conferring resistance to alpha-cypermethrin in *Anopheles funestus*

**DOI:** 10.1101/2025.04.16.649184

**Authors:** Talal AL-Yazeedi, Grâce Djuifo, Leon Mugenzi, Abdullahi Muhammad, Jack Hearn, Charles S. Wondji

## Abstract

The heavy reliance on pyrethroid-based interventions has accelerated the spread of resistance malaria vectors including *Anopheles funestus*, jeopardising control efforts. The efficacy of Insecticide-based interventions, especially insecticide-treated nets (ITNs), the cornerstone of malaria control and management, is threatened by the widespread resistance complicating malaria control. Alpha-cypermethrin, a type II pyrethroid, is increasingly utilised in various ITN formulations, including those combined with piperonyl butoxide (PBO) and chlorfenapyr-based Interceptor® G2 (IG2) nets, to enhance effectiveness against resistant mosquito populations. Therefore, understanding the molecular basis of resistance is essential to monitor and track resistance trends for an effective malaria control program. In this study, we identified a 1.4 Mb QTL on the telomeric end of the left arm of chromosome 2, conferring resistance to α-cypermethrin (*rap1* QTL). Different crossing schemes and sequencing approaches were explored to determine the most effective strategy. Individual-based QTL mapping performed on segregating individuals from an isofemale family identified a QTL at the F_7_ generation. Higher recombination density relative to the physical genome in the F_7_ isofemale family, with a recombination every 240 kb, facilitated the detection of a QTL compared to the F_2_ family (335 kb/cM). Additionally, we exploited bulk segregate analysis (BSA) between susceptible and resistant phenotypes from the F_7_ isofemale family and an F_7_ mixed cross-family to perform cost-effective and rapid QTL-mapping discovery. The strongest signal in both independent BSA analyses overlaps with the *rap1* QTL, further supporting its role in α-cypermethrin resistance. The known resistant alleles of the cytochrome P450 *CYP6P9a* and 6.5-kb structural variant within the *rap1* QTL strongly correlate with survival to α-cypermethrin. In this study, we validated that previously developed DNA-based assays, originally designed to monitor permethrin resistance, are effective for tracking resistance to α-cypermethrin as well. Additionally, we identified candidate variants that can serve as reliable markers for monitoring α-cypermethrin resistance.

**Author Summary:** In the study we used genetic crosses between resistant and susceptible *Anopleles funestus* colonies to identify quantitative trait loci (QTL) associated with the resistance to α-cypermethrin. We have identified a QTL on the left arm of chromosome 2 conferring resistance to α-cypermethrin (*rap1* QTL). Different crossing schemes and sequencing approaches were explored to determine the most effective and cost-efficient strategy to map candidate loci associated with resistance. In this study we assessed the efficiency and the cost-effectiveness of Bulk segregant analysis (BSA) mapping to detect candidate loci associated with resistance. BSA analysis enabled the detection of polymorphism in candidate regions that could serve as potential SNP-based marker to track resistance to α-cypermethrin. Previously developed SNP-based markers to track resistance to permethrin in known resistant alleles show a strongly correlation with survival to α-cypermethrin, suggesting shared mechanisms my underly resistance to both type I (permethrin) and type II (α-cypermethrin) pyrethroids.

## Introduction

Resistance to several insecticides, including pyrethroids, has been confirmed in *Anopheles funestus*, a key malaria vector across sub-Saharan Africa, jeopardising malaria control efforts. Malaria remains a global challenge, with an estimated 249 million malaria cases and 631,000 deaths recorded in 2022 alone [1]. The burden is particularly high in sub-Saharan Africa, accounting for 96% of malaria deaths globally. Insecticide-based interventions, particularly insecticide-treated nets (ITNs), have been instrumental in reducing malaria morbidity and mortality globally [2]. The WHO estimates that between 2000 and 2022, ITNs have averted an estimated 2.1 billion malaria cases, and 11.7 million malaria deaths [1].

However, the overuse of pyrethroid-based interventions has driven widespread resistance in *An. funestus*, threatening the efficacy of ITNs [3–8]. Until recently, pyrethroid-only insecticide-treated nets (pITNs), such as those containing deltamethrin, permethrin, or α-cypermethrin, were the primary interventions recommended by WHO. Heavy reliance on these insecticides has accelerated the spread of resistance, diminishing the effectiveness of pITNs and complicating malaria control efforts [9]. In response, the WHO has now endorsed next-generation insecticide-treated nets (ngITNs), which combine pyrethroids with either a second insecticide or a synergist that restores the susceptibility to pyrethroid [1]. Notable ngITNs include α-cypermethrin + chlorfenapyr (CFP) nets (Interceptor® G2), permethrin + piperonyl butoxide (PBO) nets (Olyset Plus®) and α-cypermethrin + pyriproxyfen (PPF) nets (Royal Guard®) [1, 10]. These innovative nets provide a dual mechanism with a complementary mode of action to pyrethroids: PBO restores susceptibility to pyrethroids by inhibiting the detoxification effect of cytochrome P450, chlorfenapyr is a second insecticide with a different mode of action targeting the mitochondria of insects, and pyriproxyfen is a hormone analogue that sterilises adult mosquitoes and reduces their life span [11]. With these new insecticide interventions deployed, it is critical to study the molecular basis of resistance to monitor and track resistance trends effectively. In this study, we investigated the molecular basis of resistance to α-cypermethrin, a primary pyrethroids insecticide in ngITNs, through quantitative trait loci (QTL) mapping using a segregating population from a cross between susceptible and resistant *An. funestus* laboratory colonies.

The previous identification of the rp1 QTL at the telomeric end of chromosome 2R in *An. funestus* was a significant breakthrough in understanding permethrin resistance [12]. This discovery was achieved by quantitative trait loci (QTL) using segregating individuals derived from a cross between the FUMOZ-resistant colony, originally from Mozambique, and the FANG-susceptible colony, originally from Angola [12].

Transcriptomic profiling of the FUMOZ-resistant strain revealed the over-expression of the rp1 locus duplicated CYP6P9a/b cytochrome p450 [13], which is driven by a mutation in cis-regulatory regions of both genes [4, 5, 14]. Further investigation demonstrated that a 6.5-kb intergenic structural variant between *CYP6P9a* and *CYP6P9b* enhances P450-mediated resistance, compromising the efficacy of permethrin-only bednets [8]. This molecular insight paved the way for developing a DNA-based PCR assay that allowed the detection and tracking of permethrin resistance in the field, offering a valuable tool for assessing the efficacy of permethrin-only bed nets in malaria control programs. However, these resistant markers only explain the molecular basis of resistance in southern Africa [4, 5]. The molecular basis of resistance is highly diverse and complex across Africa, with different metabolic resistance profiles, particularly cytochrome P450-mediated resistance emerging from various origins and potentially merging as they spread [15, 16]. The extensive genetic variation between regions is a major challenge preventing the design of effective resistance control and management strategies across Africa [16, 17]. However, transcriptomic analyses and large-scale genome sequencing of field-resistant populations enabled the identification of key resistance genes in *An. funestus* from different regions across Africa [4, 18]. Consequently, DNA-based assays have been successfully developed to monitor and track resistance, improving the precision of resistance management in these areas.

Previously, *Anopheles* segregating populations derived from a cross between resistant and susceptible strains were successfully used to map loci conferring resistance to pyrethroids. These experiment designs heavily relied on laboratory colonies where clear segregating of susceptibility and resistance phenotype was possible, and the strains were compatible for crossing. In this study, we leveraged a segregating population derived from a cross between the resistant FUMOZ and susceptible FANG colonies to uncover the molecular basis of α-cypermethrin resistance. We integrated different sequencing techniques and crossing strategies to determine the most cost-effective approach to map resistant loci. To reduce the cost of sequencing while achieving sufficient genome-wide genotyping of segregating individuals, we employed double digest restriction-site associated DNA sequencing (ddRADseq) on an isofemale family segregating population at the F_2_ and the F_7_ generation. However, mapping resistant loci from segregating individuals derived from isofemale families using traditional methods is time-consuming and challenging in the field where there is little to no segregation in susceptibility phenotype from the same geographical region. Therefore, we exploited a strategy of pooling DNA from individuals with extreme phenotypes to perform a cost-effective, rapid, and efficient QTL mapping discovery. Next-generation sequencing coupled with bulk segregate analysis (NGS-BSA) was performed on pools of Dead and Alive, segregating individuals from the F_7_ iso-female family and a mixed cross family independently.

This design offers a valuable, rapid and cost-efficient model to monitor resistance in *Anopheles* species, especially in scenarios where segregating susceptibility is no longer prevalent due to widespread resistance and resistance to new insecticide formations is rapidly spreading.

We observed higher recombination rates in the F_7_ generation, which expanded the genetic map and facilitated the detection of the *rap1* QTL compared to the F_2_ generation. Genotyping with molecular markers from the *rap1* QTL revealed a strong association between the resistant copy of the *rap1* locus and α-cypermethrin resistance. BSA offers a fast and cheap approach to map loci associated with resistance from isofemale and mixed cross families. Integrating BSA with the segregating individuals-based QTL mapping, we were able to investigate polymorphism within the *rap1* QTL and determine candidate variants for molecular diagnostic susceptibility assay. This study provides a framework for future studies aimed at mapping resistant loci to already deployed insecticides or rapidly evolving resistance to new insecticide formulations

## Results

### Higher recombination rate in F_7_ Iso-female family revealed by Genetic Linkage Mapping (GLM)

Genetic linkage maps (GLM) were constructed for an F_2_ and an F_7_ generation of *An. funestus* segregating population to investigate the *An. funestus* recombination landscape and scan for individual-based quantitative trait locus (QTL) associated with α-cypermethrin resistance. A FUMOZ iso-female mated with a FANG male initiated the F_2_ and the F_7_ segregating populations, named the F_2_ and the F_7_ families respectively (see methods for details). Markers from the F_2_ family and the F_7_ family genotyped using double digest restriction-site associated DNA sequencing (ddRADseq) were separated into three linkage groups corresponding to *An. funestus* chromosomes based on the pairwise recombination fraction (RF) between markers and LOD score (Figure 1A, 1B and S1).

**Figure 1.**
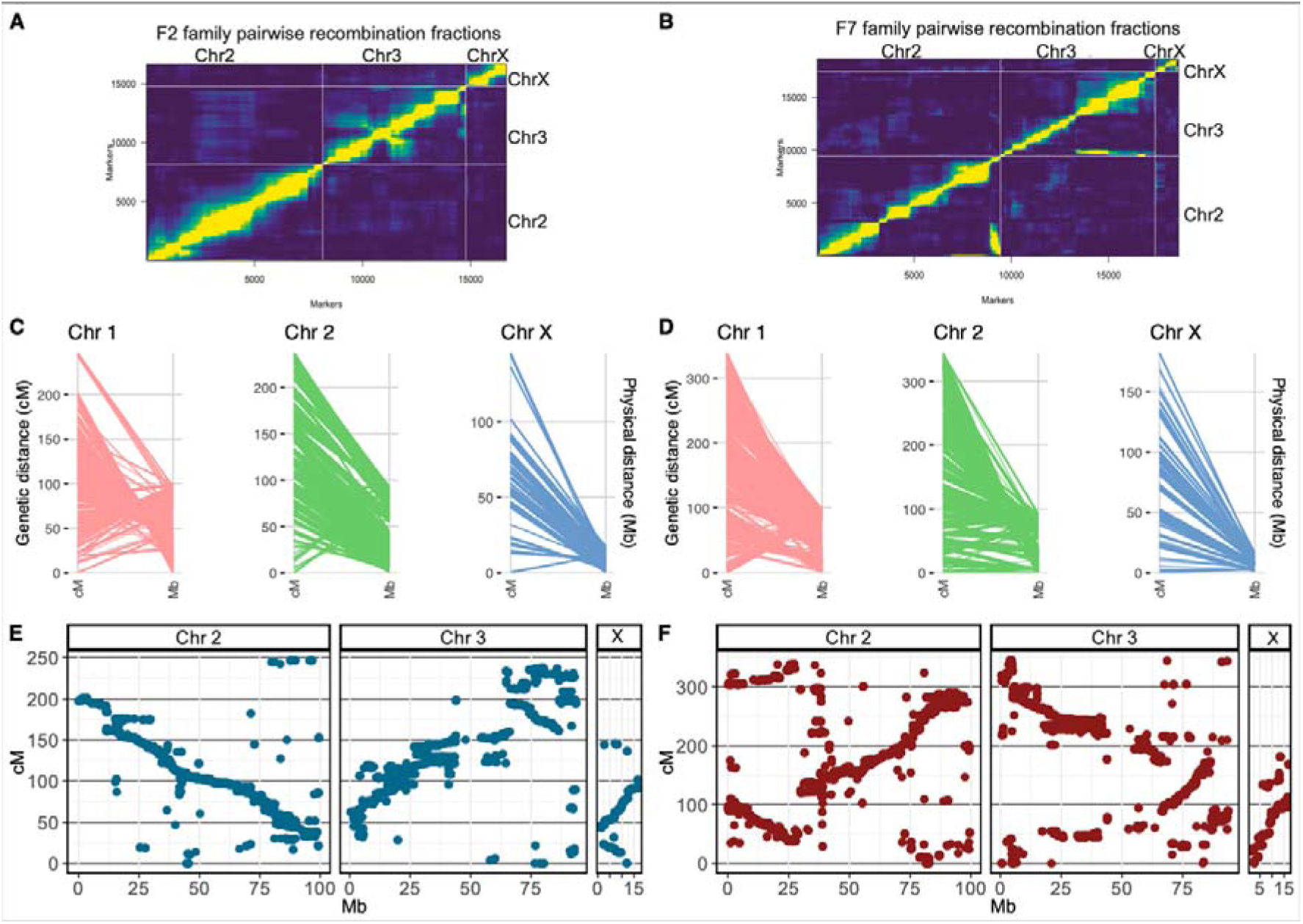
Genetic linkage maps and their relationship with the FUMOZ genome physical distance. Markers’ pairwise recombination fraction and LOD score for all markers pairs in the F_2_ (**A**) and F_7_ (**B**) iso-female family illustrate a strong linkage between markers in each separate linkage group. Markers were separated into three main linkage groups corresponding to chromosomes. Plot of estimated recombination fractions (upper-left triangle) and LOD scores (lower-right triangle) for all pairs of markers. Yellow indicates linked (large LOD score or small recombination fraction), and blue indicates no linkage (small LOD score or large recombination fraction). Markers’ genetic linkage distance in (cM) and their corresponding location in the FUMOZ genome physical distance (Mb) are highlighted for the F_2_ (**C**) and the F_7_ (**D**) iso-female family. The Marey plots illustrate the relationship between F_2_ (**A**) and F_7_ (**B**) families’ cumulative genetic distance in cM plotted against the FUMOZ genome physical distance. Inversions in the genetic linkage map compared to the FUMOZ genome physical distance are highlighted by the inverse relationship between the recombination distance in cM and the physical distance in Mb, with regions of high genetic distance corresponding to short physical distance.

The overall GLM length in the F_7_ family (865.5 cM) is larger than the F_2_ family (629.7 cM) due to the increased number of recombination events by multiple successive generations of sibling mating (Table 1) (Figure 1). We identified a higher recombination density in the F_7_ family of 4.1 cM/Mb (corresponding to a ratio of physical and genetic distance genome size of 240 kb/cM). This contrasts with the F_2_ family of 2.1 cM/Mb (corresponding to 335 kb/cM). In both families, recombination was higher in the X chromosome compared to the autosomes, with recombination events occurring in 100 kb and 120 kb in the F_7_ and the F_2_ family, respectively (Table 1). In both families, the ratio of heterozygote markers in the linkage map is higher than the homozygous markers almost reflecting the expected genotype ratio of an F_2_ segregating population (Table 1) (Figure S2). Parental background for markers at each position on the genome for all individuals and the ratio of markers in each genetic linkage map is illustrated in supplementary Figures S3, S4, S5 and S6.

**Table 1.**
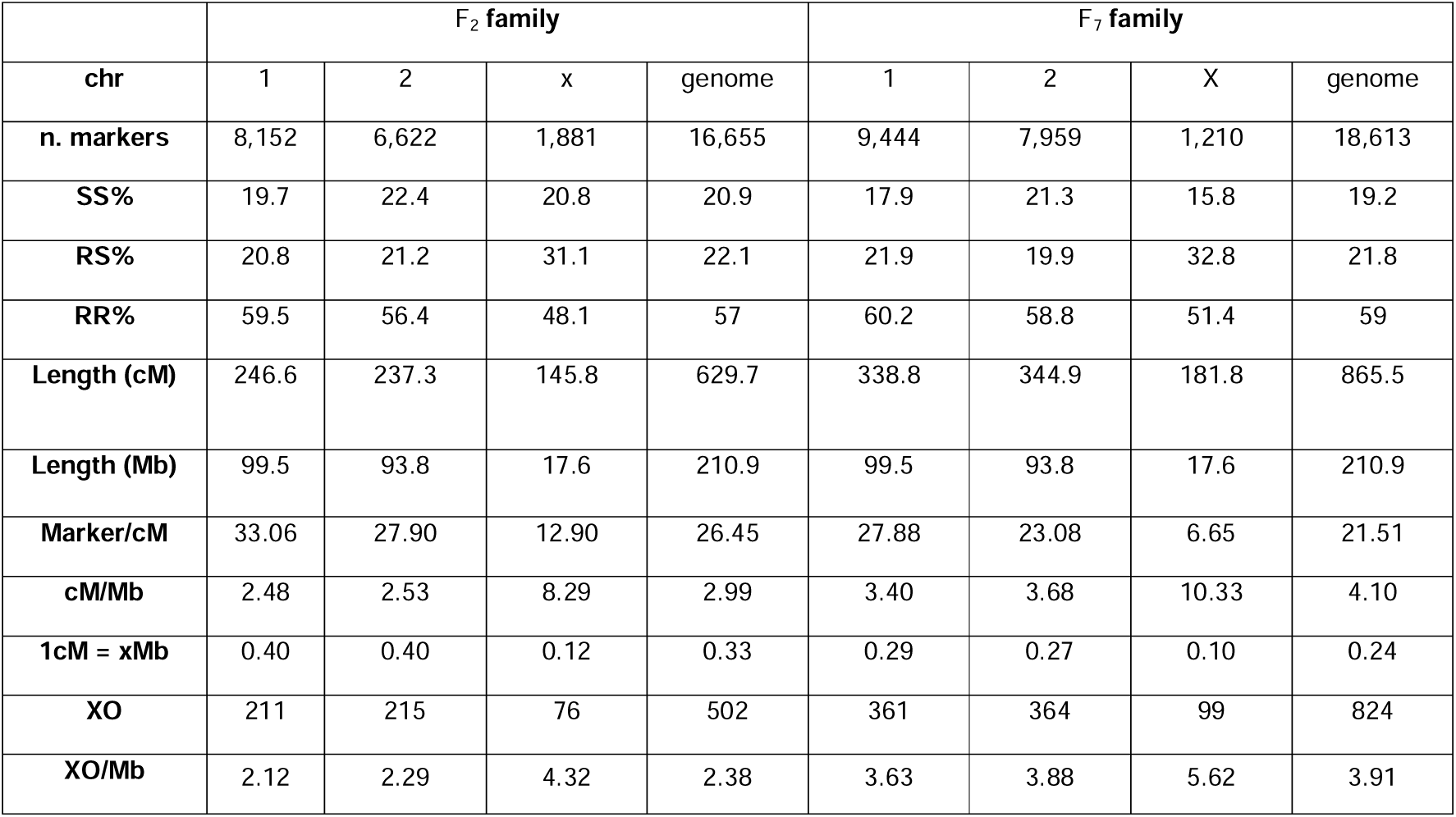
F_2_ and F_7_ family genetic linkage map metrics. The table highlights the number of markers in each of the F_2_ and the F_7_ family (n.markers), the ratio of parental genotypes in the genetic linkage maps, the length of the map in centimorgan (Length cM), the number of markers in 1cM (Markers/cM), the recombination rate of cM/Mb, number of crossovers (XO) and number of crossovers per Mb (XO/Mb).

Markers with severe segregation distortion were removed from the linkage map, while markers with segregation distortion p-value greater than the family-wide Bonferroni adjusted alpha level of (0.05/no of markers) were highlighted (Figure S2). Marey plots for the F_2_ and the F_7_ family illustrate the relationship between the physical distance in Mb of the FUMOZ reference genome and the recombination distance in cM for each chromosome (Figure 1 E and F). The non-linear relationship manifested by the Marey plots illustrates the structural difference between the FANG and the FUMOZ genomes (Figures 1C and D).

Investigation of recombination hotspots identified higher crossover events in the F_7_ family with a total of 824 (corresponding to a ratio of 3.91 crossover events per Mb). This contrasts with the F_2_ family of a genome total of 502 crossover events (corresponding to 2.38 crossovers per Mb) (Table 1) (Figure 2 A and B). The relationship between markers’ recombination rate per Mb (cM/Mb) and physical length in Mb in both F_7_ and F_2_ families was fitted with a multiplicative inverse function, illustrating the inverse relationship between recombination rate and physical distance. The recombination rate is high when the physical distance between markers is small and decreases as the physical distance between markers increases when they are further apart on the chromosome (Figure 2 C and D). To highlight areas of dense and low recombination in the F_2_ and the F_7_ family compared to the FUMOZ reference genome, recombination rates (cM/Mb) were computed by dividing genetic distance into physical distances in a window size of 1LMb (Figure 2 E and F).

**Figure 2.**
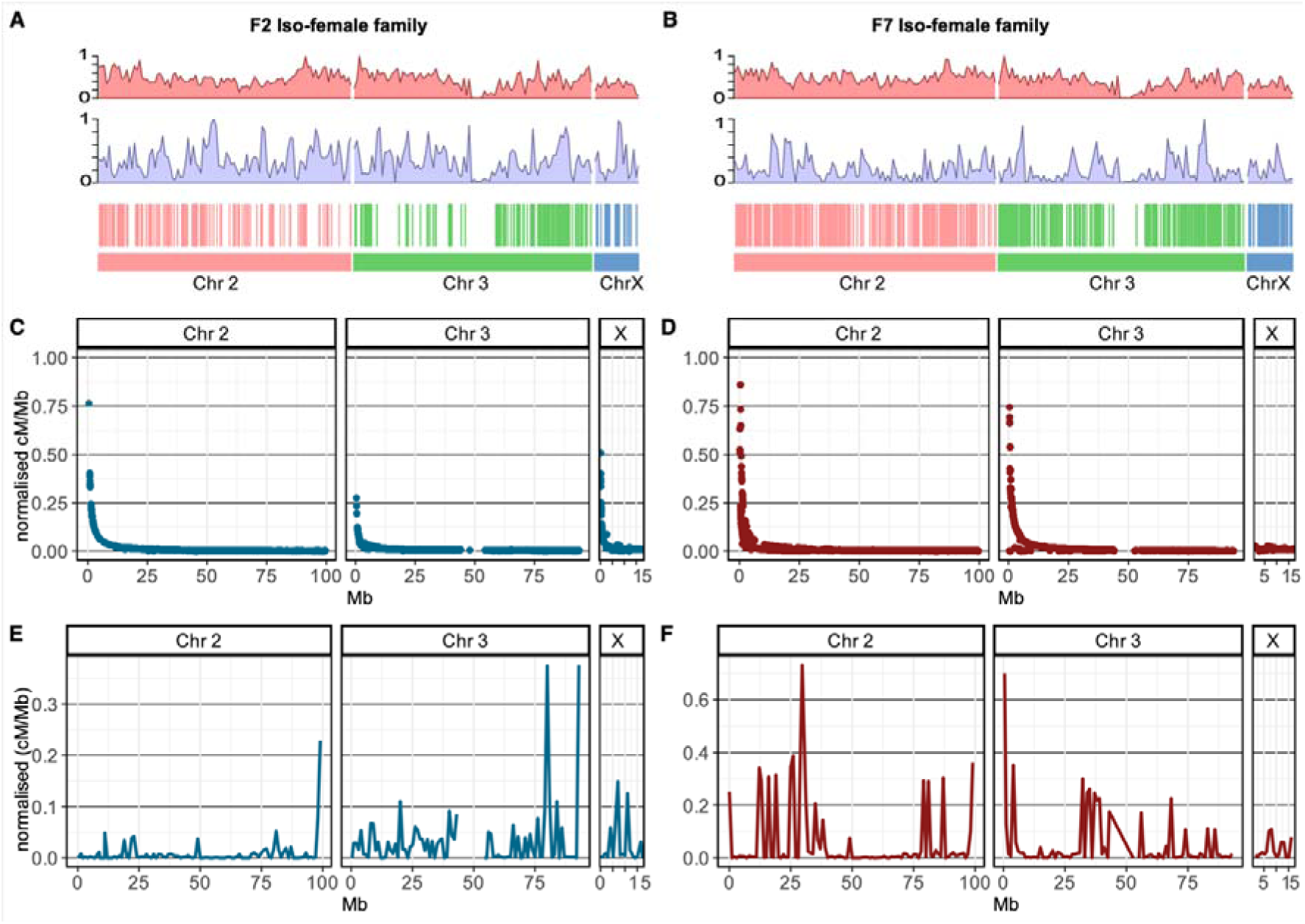
Recombination hotspots and recombination rate in the F_2_ and F_7_ iso-female families. The number of crossovers in the F_2_ (**A**) and the F_7_ (**B**) genetic linkage maps were calculated, and their relative positions in the FUMOZ reference genome are highlighted in vertical stripes with a similar colour to the corresponding chromosome. The number of crossovers varies along the chromosome with limited crossover events around the centromere; this is more evident in Chromosome 3. Density plots of monomorphic (purple) and polymorphic (salmon) variants for each chromosome are highlighted in density plots. In total, 59,161 variants were identified to be polymorphic in the F_7_ family, and 5,119 variants were monomorphic. In the F_2_ family, 46,160 variants were polymorphic and 9,360 were monomorphic. The recombination rate for each maker was calculated by dividing the genetic distance (cM) by the physical distance (Mb) (cM/Mb) and then normalised by dividing by the total recombination distance (F_2_ = 629.7 and F_7_ = 865.5). This normalized rate for the F_2_ (**C**) and F_7_ (**D**) families was plotted against the physical distance in Mb on the x-axis. The plot reflects the inverse relationship between the recombination rate and physical distance where a high recombination rate occurs over a small physical distance and the rate decreases as the physical distance increases. Areas of dense and low recombination in the F_2_ (**E**) and the F_7_ (**F**) families were highlighted by computing recombination rate (cM/Mb) in a window size of 1Mb (y-axis) and visualised against the FUMOZ genome physical distance (x-axis).

### Individual-based QTL scan identified a major QTL responsible for **α**-cypermethrin on the left arm of chromosome 2

Individual-based QTL scan using Haley-Knott regression with the resistance to α-cypermethrin phenotype coded in a binary format (resistant =1, and susceptible = 0) detected a major QTL in the F_7_ family conferring resistance to α-cypermethrin (Figure 3A). On the contrary, no QTL peaks were discovered in the F_2_ family that exceeded the genome-wide threshold (Figure 3B).

**Figure 3.**
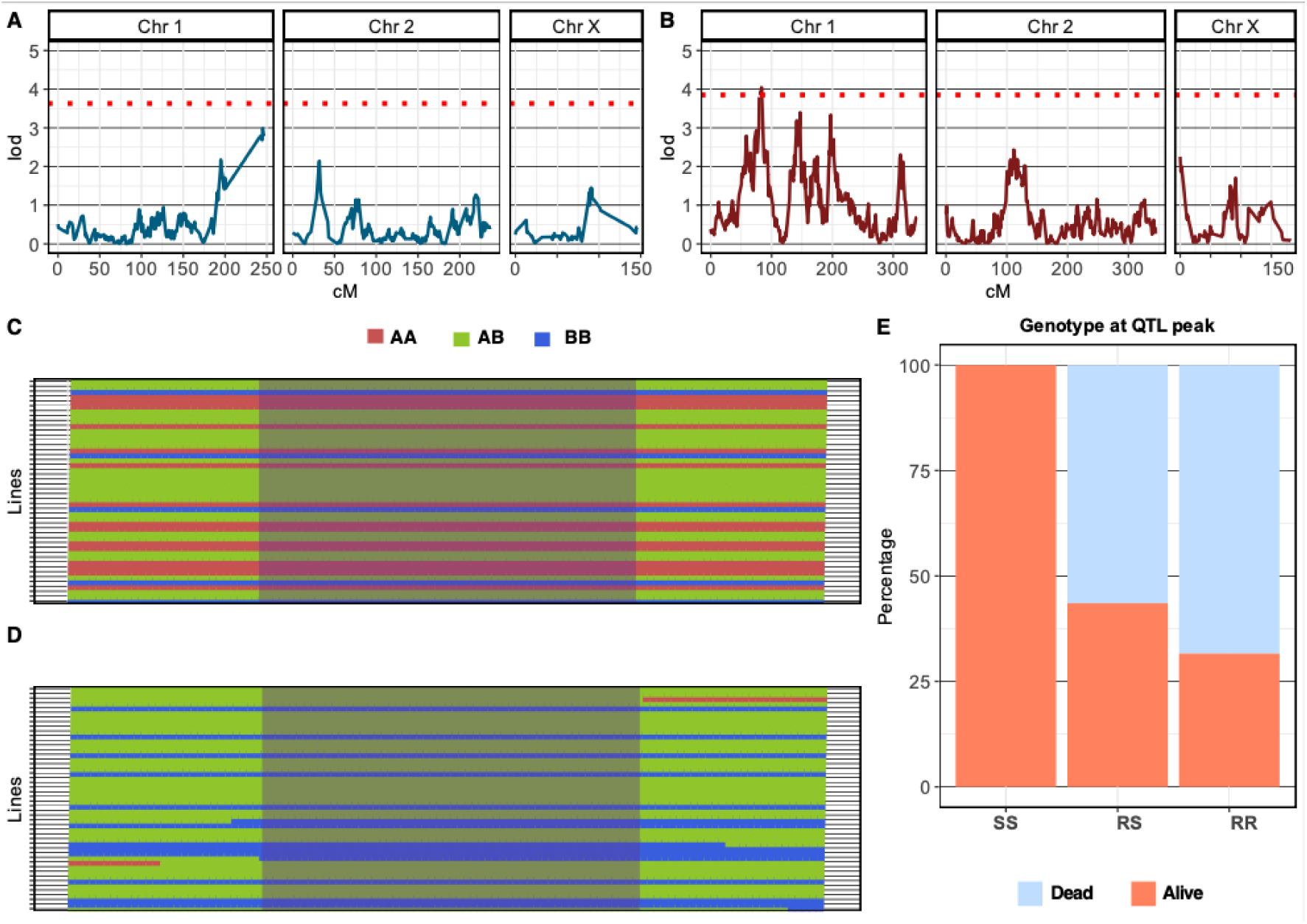
QTL mapping and the distribution of resistant phenotype at the QTL peak. To detect a QTL associated with α-cypermethrin resistance in the F_2_ (**A**) and the F_7_ (**B**) families a QTL scan in a single mode was conducted using Haley-Knott regression with the resistance to α-cypermethrin phenotype coded in a binary format (resistant =1, and susceptible = 0). The QTL in the F_7_ family (**B**) was determined based on the genome-wide threshold (red-dotted line) corresponding to an LOD score of 5.31. QTL markers Genotypes from individuals of the F_7_ isofemale family on the y-axis plotted along the (QTL) genetic distance (79.5555 to 83.3190 cM) reveal a difference in the parental genotype composition (AA FANG and BB FUMOZ) between dead (**C**) and alive (**D**) individuals. The QTL peak highlighted in purple indicates no alive individuals have (SS) FANG genotype at the QTL peak. Genotype percentage at the QTL peak indicates that all individuals with susceptible (SS) genotypes were susceptible to α-cypermethrin. Meanwhile, (65%) of individuals with the resistant genotype (RR) at the QTL peak were resistant and (56.45%) of individuals with the heterozygote genotype (RS) were resistant to α-cypermethrin (**E**).

The identified resistance to α-cypermethrin QTL is ∼ 1.44 Mb, equivalent to 3.23 cM in the F_7_ family genetic linkage map (Figure 3). We named this QTL resistance to α-cypermethrin QTL (*rap1*) in accordance with the previously named QTL identified to confer resistance to permethrin (*rp1* QTL) [12, 14]. The QTL was determined based on a genome-wide p-value threshold < 0.05 corresponding to a LOD threshold of 3.82 obtained through 1,000 times permutation of the QTL scan (Figure 3B). The QTL was detected in cM using genetic linkage distance from the F_7_ family genetic linkage map in the distance from 79.5555 cM to 83.3190 cM corresponding to two regions on chr 2 from 4,558,633 bp to 4,558,748 bp (155 bp) and from 8,127,007 bp to 9,567,884 bp (1,440,877 bp). The *rap1* QTL represents 0.5% of the total genome size of the FUMOZ reference genome.

The peak of the QTL corresponds to a LOD score of 5.31 located at a genetic linkage position of 81.2 cM on Chr2. The QTL peak corresponds to a physical position starting at 8,194,020 bp and ending at 9,286,436 bp (1,092,416 bp) on Chr2 (Figures 3C and 3D). The QTL is estimated to explain 22.5% variance in the binary resistant phenotype from 96 individuals using a regression model (formula: y ∼ QTL, p-value (Chi2): 4.85e-06). We analysed the distribution of genotypes between dead and alive at a marker close to or at the QTL peak of 81.2 cM. We found a strong association between individuals with the resistant genotype at that marker and survival to α-cypermethrin (Chi^2^ = 19.212, p-value = 6.733e-05). All 15 individuals with the FANG phenotype (AA) at the QTL peak were susceptible to α-cypermethrin (Figure 3E). On the other hand, 13/20 (65%) of individuals with homozygotes FUMOZ genotype (BB) at the QTL peak were resistant while 6/20 (35%) were susceptible. From the total individuals with heterozygote phenotype (BB) at the QTL peak 35/62 (56.45%) were resistant and 27/62 (43.54%) were susceptible to α-cypermethrin (Figure 3E and S7).

There are 113 annotated genes overlapping the QTL region from 4,558,633 bp to 4,558,748 bp and from 8,127,007 bp to 9,567,884 bp on Chr2 from the FUMOZ reference genome (Dataset 1). Gene ontology analysis of genes in the *rap1* QTL region identified 16 genes with oxidoreductase activities (GO:0032934) including 11 *CYP450* genes with a monooxygenase activity (GO:0004497) all from the CYP6 cluster. Notably, *CYP6P9a* and *CYP69b* previously were demonstrated to confer resistance to permethrin, deltamethrin and carbamate [5, 8, 19]. Other CYP6 genes identified in this region includes *CYP6AD1*, *CYP6AA2*, *CYP6AA1*, *CYP6P2*, CYP6P1 and CYP6P5. The *rap1* QTL contains genes with carboxylic ester hydrolase activity (GO:0052689) including Carboxylic ester hydrolase AFUN015787 and AFUN015793 (Dataset 2).

In addition to detoxification genes, the QTL contains genes involved in active ion transmembrane transporter activity (GO:0022853), including sodium-calcium exchanger solute carrier family 8 (AFUN020405), sodium-potassium-transporting ATPase subunit alpha (AFUN008354), chloride channel activity (GO:0005254) glutamate receptor ion channel gene (AFUN020159), and voltage-gated chloride channel gene (AFUN008007).

Overlapping SNPs markers at the QTL peak with highest LOD score with gene regions identified overlap with genes potentially associated with resistance including sodium-potassium-transporting ATPase subunit alpha (AFUN008354), sodium-calcium exchanger solute carrier family 8 (AFUN020405), *CYP6AA1* and *CYP6P2* (Dataset 3). Investigating the underlying function of these SNPs revealed missense SNPs in *CYP6AA1* and *CYP6AA1* (Dataset 3).

### Correlation between genotypes at *CYP6P9a* and the 6.5kb structural variant and **α**-cypermethrin resistance

Genotyping the dead and alive individuals from the isofemale F_7_ family using molecular markers revealed that *An. funestus* with a homozygous form of the structural variant (SV+/SV+) or homozygous *CYP6P9a* resistant (RR) genotypes, whether inherited independently or together, have an increased advantage to survive exposure to α-cypermethrin. A significant difference was observed in the distribution of the 6.5kb (SV+/SV+) between dead (15.8%) and alive (84.2%) *An. funestus* from the F_7_ isofemale family after exposure to α-cypermethrin (Chi^2^=25.53, p-value = 2.854e-06) (Table S2). Reciprocally, the absence of the 6.5kb SV (SV-/SV-) was greater in the dead (95%) compared to the alive (5%). *An. funestus* homozygous individuals for the 6.5kb have an odds ratio of 101.3 more likely to survive when exposed to α-cypermethrin compared to homozygous without the structural variant (SV-/SV-) (Estimation of odds ratio OR = 101.33, 95% confidence interval CI = 9.58 - 1072.04, P value = 0.0001), and 4.47 time more likely to survive compared to individuals heterozygote to the 6.5kb SV (SV+/SV-) (OR = 4.47, 95% CI = 1.17-17.06. P value = 0.028) (Table S3). Additionally, Individuals who are heterozygote for the 6.5kb SV (SV+/SV-) are 22.65 times more likely to survive exposure to α-cypermethrin compared to homozygous individuals (SV-/SV-) (OR = 22.65, 95% CI = 2.84 – 180. 85, P value = 0.0032) (Figure 4) (Dataset 4).

**Figure 4.**
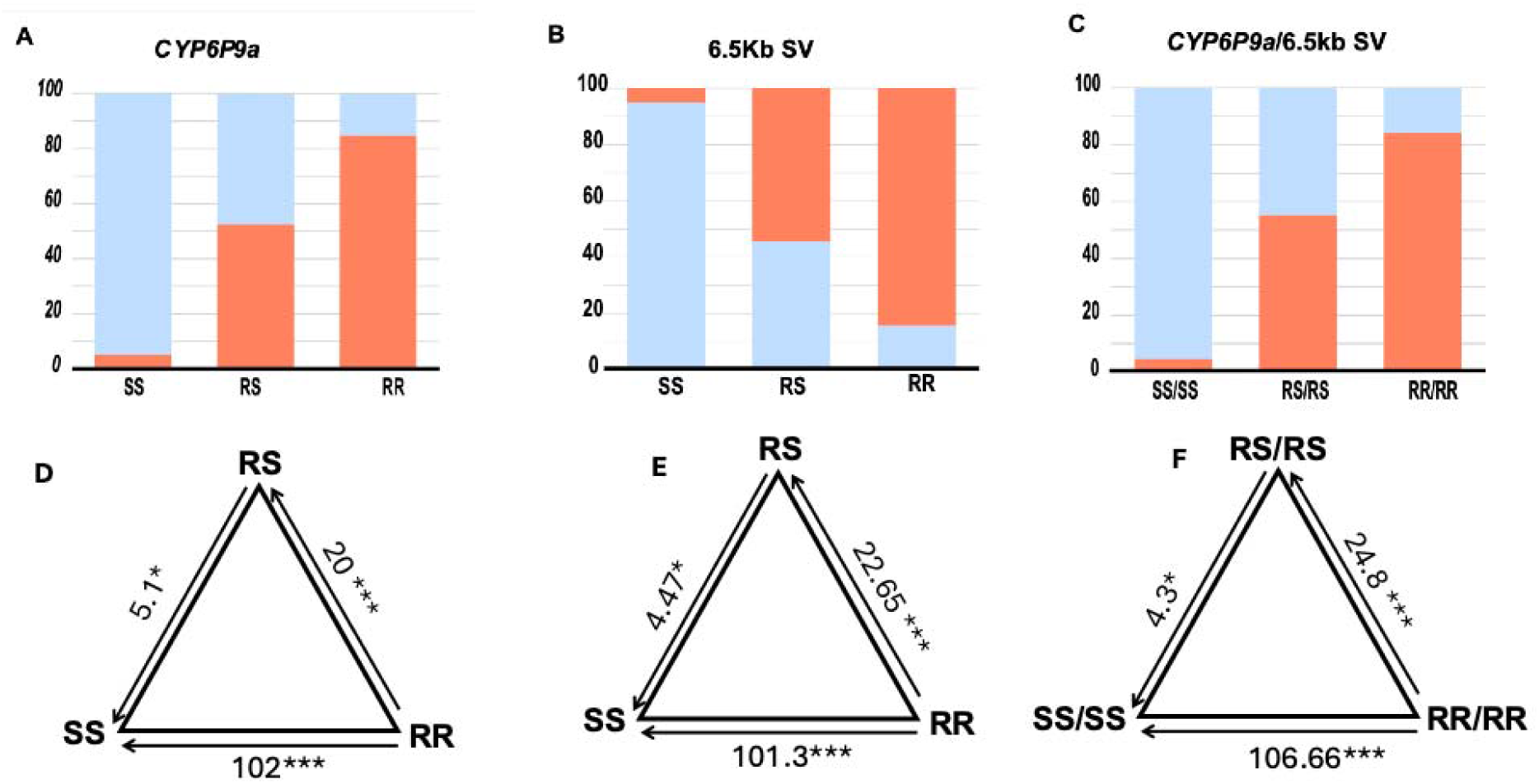
Correlation between *CYP6P9a* and the 6.5kb structural variant and α-cypermethrin resistance. Distribution of *CYP6P9a* (**A**) and 6.5Kb insertion (**B**) independently and together (**C**) between dead and alive segregating individuals from the F_7_ isofemale family indicates the individuals with the resistant allele are more likely to survive exposure to α-cypermethrin. Estimation of odds ratio (OR) indicates that individuals with homozygote resistant allele of *CYP6P9a* are 102 more likely to survive exposure to α-cypermethrin compared to individuals with the susceptible allele (**D**). Individuals with the 6.5kb insertion are 101.3 more likely to survive compared to individuals without the insertion (**E**). Individuals with the resistant copy of CYP6P9a and the 6.5kb insertion are 106.66 times more likely to survive exposure to α-cypermethrin compared to individuals’ homozygote for the susceptible copy of *CYP6P9a* and lack the 6.5kb insertion (**F**). Odds ratios are given with asterisks indicating the level of significance.

Moreover, Individuals with homozygote *CYP6P9a* resistant (RR) genotype are strongly associated with α-cypermethrin resistance. There is a strong association between individuals with *CYP6P9a* resistant (RR) genotype (85%) and survival to exposure to α-cypermethrin (Chi^2^ = 23.665, p-value = 7.264e-06). Individuals with a resistant (RR) genotype (85%) are 102 more likely to survive exposure to α-cypermethrin compared to individuals homozygous to the susceptible form of *CYP6P9a* (SS) (OR = 102, CI = 9.64 - 1078.43, Pvalue = 0.0001), and 5.1 times more likely to survive compared to the (RS) (OR = 5.1, 95% CI = 1.34 – 19.34, P value = 0.016) (Table S2). Individuals with one resistant copy of *CYP6P9a* (RS) are more likely to survive exposure to α-cypermethrin with an odds ratio of 20 compared to homozygous individuals for the susceptible form of *CYP6P9a* (SS) (95% CI = 2.49 – 160.04, P value = 00048) (Figure 4) (Dataset 5).

We investigated the combined effect of the 6.5kb insertion and the homozygote-resistant copy of the CYP6P9a (SV+/SV+/RR) on α-cypermethrin resistance. There is a significant association between individuals with combined homozygote copies of the resistant form of the 6.5kb and cyp6p9a (SV+/SV+/RR) and resistance to α-cypermethrin (Chi^2^ = 26.728, p-value = 1.571e-06), where 84.2% were resistant and 15.8% were susceptible to α-cypermethrin. Double homozygous individuals with one resistant copy of 6.5kb insertion and cyp6p9a are more likely to survive exposure to α-cypermethrin compared to double homozygote to the susceptible alleles (SV-/SV-/RR) with an odd ratio of 106.6 (95% CI 10.1 – 1126.04, P value = 0.0001), and more likely to survive compared to double heterozygote with odd ratio of 4.3 (95% CI = 1.1 – 16.4, P value = 0.03). Additionally, double heterozygote individuals (SV+/SV-/RS) are 24.8 times more likely to survive exposure to α-cypermethrin compared to double homozygote to the susceptible copies (SV-/SV-/SS) (Figure 4) (Dataset 5) (Table S2 and S3).

### BSA-identified candidate regions associated with **α**-cypermethrin resistance

Next-generation sequencing coupled with bulk segregate analysis (NGS-BSA) exploits a strategy of pooling DNA from individuals with extreme phenotypes to perform a cost-effective, rapid, and efficient QTL mapping. This method was used successfully to correlate phenotype with genotype and identify candidate regions from whole genome sequencing of insect DNA pools [20, 21]. To assess the effectiveness of the NGS-BSA approach in identifying candidate regions associated with α-cypermethrin resistance in segregating *An. funestus* population two genomic pools were prepared one for resistant individuals (Alive after 1h exposure to α-cypermethrin) and one for susceptible individuals (Dead after 1h exposure to α-cypermethrin) from two different crossing strategies. 1) from the same F_7_ generation iso-female family used for individual-based QTL analysis and 2) from an F_7_ generation segregating population produced from mixed crossings (see method for details). Different methods were employed to identify candidate regions and evaluate the statistical significance of the identified candidate regions, including the QTL-seq method and the G’ method [22–24].

BSA using the F_7_ isofemale family DNA pools identified a ∼ 14.57 Mb candidate region associated with α-cypermethrin resistance, spanning from 183 bp to 14,572,176 bp overlapping with 1,120 genes, on the left arm of Chr2 constituting 6.9% of the FUMOZ reference genome (Figure 5) (Table 2 and Table S4). The individual-based QTL identified from the same segregating individuals of the F_7_ isofemale family (*rap1* QTL) is located within the candidate region identified by BSA. However, the difference in size between the detected individual-based rap1 QTL and the BSA analysis candidate region is 13,131,001 bp. We detected 39,512 significant SNPs with a p-value less than or equal to the False Discovery Rate (FDR) threshold of alpha = 0.01. We identified 872 missense SNPs within the significant SNPs, 17 splice region SNPs, one start lost SNP and 3 stop gained SNPs (Figure S9 and Table S6). Missense SNPs with a substantial frequency difference between susceptible and resistant were detected in genes belonging to the cyp6 and the cyp32 clusters. A full list of SNPs with a significant allele frequency difference between resistant and alive bulks with the corresponding effect of these SNPs is provided in supplemental (Dataset 8).

**Figure 5.**
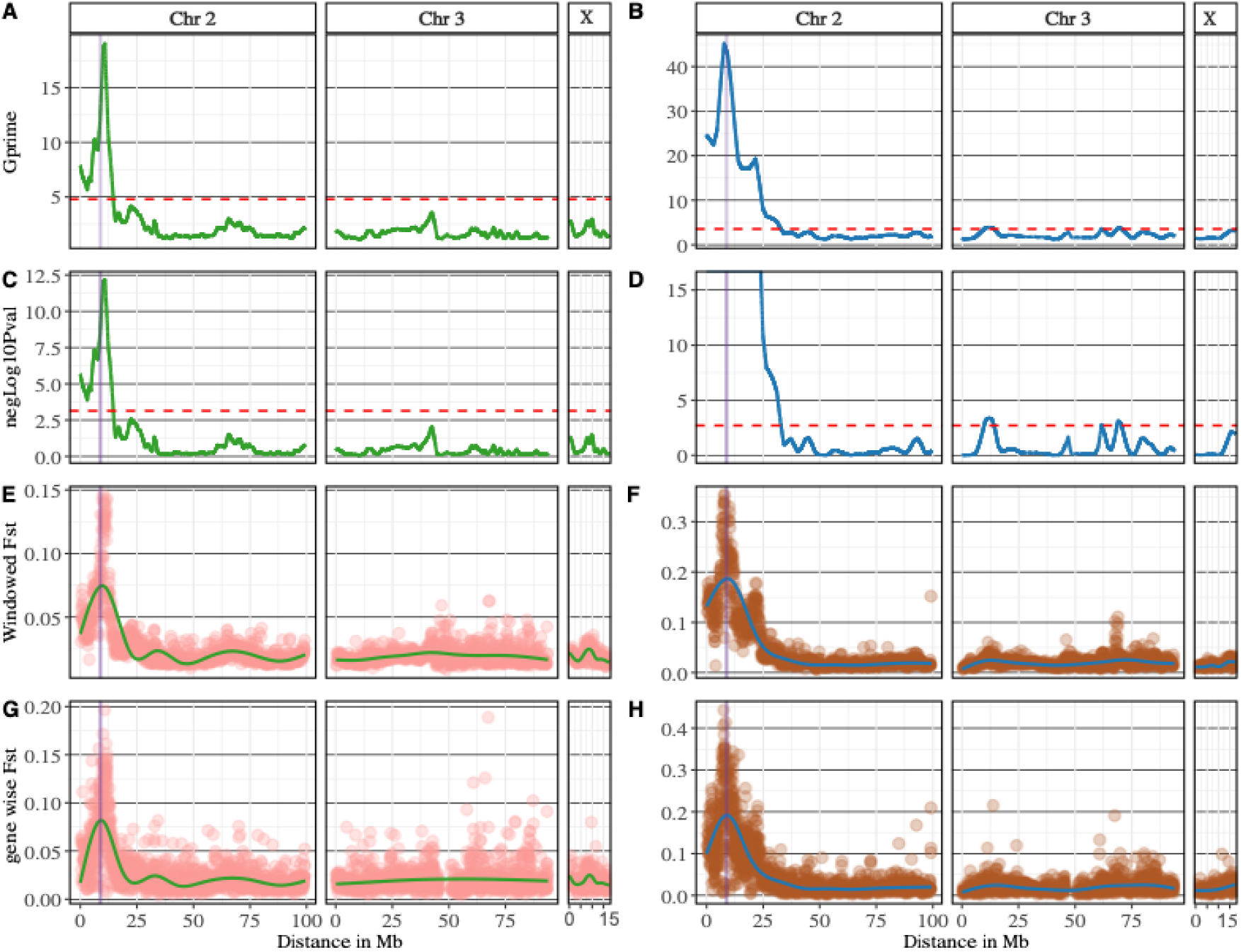
BSA analysis and genetic differentiation between dead and alive DNA bulks in F_7_ isofemale family and F_7_ mixed cross family. A tricube-smoothed G-statistics (G’) (y-axis) was calculated in a sliding window of 1 Mb across the genome distance in Mb between dead and alive bulks in the F_7_ isofemale family (**A**) and the F_7_ mixed cross-family (**B**). The *rap* QTL identified using individual base QTL is highlighted in purple in all plots. The heights G-prime (G’) peak in the F_7_ isofemale family and the mixed-cross family correlates with *rap* QTL and the red-dotted line indicates the G’ threshold at the genome-wide false discovery rate (FDR) of 0.01. A p-value was determined by comparing the G’ values to a non-parametrically estimated null distribution, with the assumption that the distribution of the G’ is a proximately log-normal. Negative log10 p-value and Benjamini-Hochberg adjusted p-values were calculated on the (y-axis) for F_7_ isofemale family (**C**) and F_7_ mixed cross family (**D**), red-dotted lined indicates the threshold at a genome-wide FDR of 0.01. The negative log10 p-value at the left arm of Chr 2 in the F_7_ mixed cross family is equal to infinity because the p-value is effectively equal to zero, indicating a highly significant result. Genetic differentiation (*Fst*) in a window size of 1Mb between dead and alive bulks in the F_7_ isofemale family (**E**) and the mixed F_7_ mixed-cross family (**F**) indicate a high genetic differentiation correlating with *rap1* QTL. Similarly, Gene-wise *Fst* between dead and alive in the F_7_ isofemale family (**G**) and the F_7_ mixed cross family (**H**) reveal a high genetic differentiation correlating with the *rap1* QTL.

**Table 2.** BSA analyses using dead and Alive DNA pools from the F_7_ isofemale family and mixed cross family.

|  | Total number of SNPs used in BSA | Number of Significant SNPs. Alpha <= 0.01 | Significant SNPs within the individual-based QTL | Synonymous variants | Missense SNPs | Detected candidate regions | Genes in candidate regions | Length of candidate region - (percentage of total genome) |
| --- | --- | --- | --- | --- | --- | --- | --- | --- |
| F<sub>7</sub> iso-female family | 551,221 | 39,512 | 5158 | 4,070 | 872 | 1 | 1,120 | 14.36 Mb - (6.9 %) of total genome |
| F<sub>7</sub> mixed cross | 582,672 | 114,108 | 4500 | 10,339 | 2,428 | 4 | 2,577 | 65.7 Mb – (31.1%) of total genome |

In the F_7_ mixed crosses family, we identified four candidate regions, a candidate region was located on Chr2 and three on Chr3 (Table 2 and S8). BSA analysis on DNA pools from mixed-cross identified significant regions with a total length of 39.7 Mb, constituting 18.67% of the genome and overlapping with 2,577 genes (Table 2 and Dataset 9). A total of 114,108 SNPs were identified as significant with a p-value less than or equal to the FDR threshold of alpha = 0.01, and 4,026 of those SNPs were identified to be missense (Table 2). The effect of significant SNPs was estimated and overlapped with corresponding genes (Dataset 10).

Additionally, BSA analysis was conducted using indels alone. In the F_7_ isofemale family, a candidate region was detected on the telomeric end of the chromosome 2 left arm from 1,5096 to 14,376,563, overlapping with the candidate region detected using SNPs BSA (Table S7 and Dataset 11). On the other hand, in the mixed-cross family, BSA analysis using indels detected seven candidate regions: one candidate region on Chromosome 2, four on Chromosome 3 and two on Chromosome X. All the detected candidate regions using SNPs-BSA in the F_7_ mixed cross family were detected using the indels-BSA (Table S8 and Dataset 12).

### BSA revealed a strong signal, polymorphism and high genetic differentiation in key detoxification SNPs within and outside the rap1 QTL region

The *rap1* QTL is located within candidate regions with the strongest signal identified by Bulk Segregant Analysis (BSA) in both the F_7_ isofemale family and mixed cross family (Figure 5 A and B). The highest peaks in candidate regions identified from BSA are physically close to the peak of the *rap1* QTL, which spans from 8,194,020 bp to 9,286,436 bp. Specifically, the peak value for the difference in SNP index between dead and alive pools from the F_7_ iso-female family is located at 9,326,280 bp, while in the F_7_ mixed cross family, the peak value for the delta-SNP index is located at 10,890,283 bp (Table S4). However, the peak for the G prime statistic occurs at different locations, at 10,880,716 in the F_7_ isofemale family bp and 7,781,510 bp in the F_7_ mixed cross family (Table S5). This alignment between the *rap1* QTL and the highest peaks detected by BSA suggests the significant role of the *rap1* QTL in conferring resistance to α-cypermethrin.

Additionally, by calculating the average *Fst* gene-wise and in a window size of 1 Mb, high genetic differentiation (*Fst)* was observed at the highest peak identified from BSA analyses (Figure 5 F and H). Further investigation of the polymorphisms within the *rap1* QTL from the BSA variants revealed missense SNPs and three prime UTR polymorphisms in detoxification genes, especially those in the cytochrome P450 *CYP6* cluster (Table S10 and Table S11). The polymorphisms at the three prime UTR are of potential significance since genes from the *CYP6* cluster, particularly *CYP6P9a* and *CYP6P9b* were highlighted to be overexpressed in the resistant-FUMOZ colony. In the F_7_ isofemale family, we identified 14 missense SNPs with significant allele frequency difference in 9 different genes within the *rap1* QTL (Table S10). In the F_7_ mixed cross family, we identified 12 missense SNPs with significant allele frequency difference in 8 different genes within the *rap1* QTL (Table S11). In total, eight missense SNPs were found to be shared between the F_7_ isofemale and mixed cross families (Figure 6 and Table S12). Notably missense SNP in the *CYP6AA1* is significantly differentiated between susceptible and resistant F_7_ isofemale and mixed cross families and was detected amongst the QTL peak SNPs in the conventional QTL mapping analysis (Figure 6).

**Figure 6.**
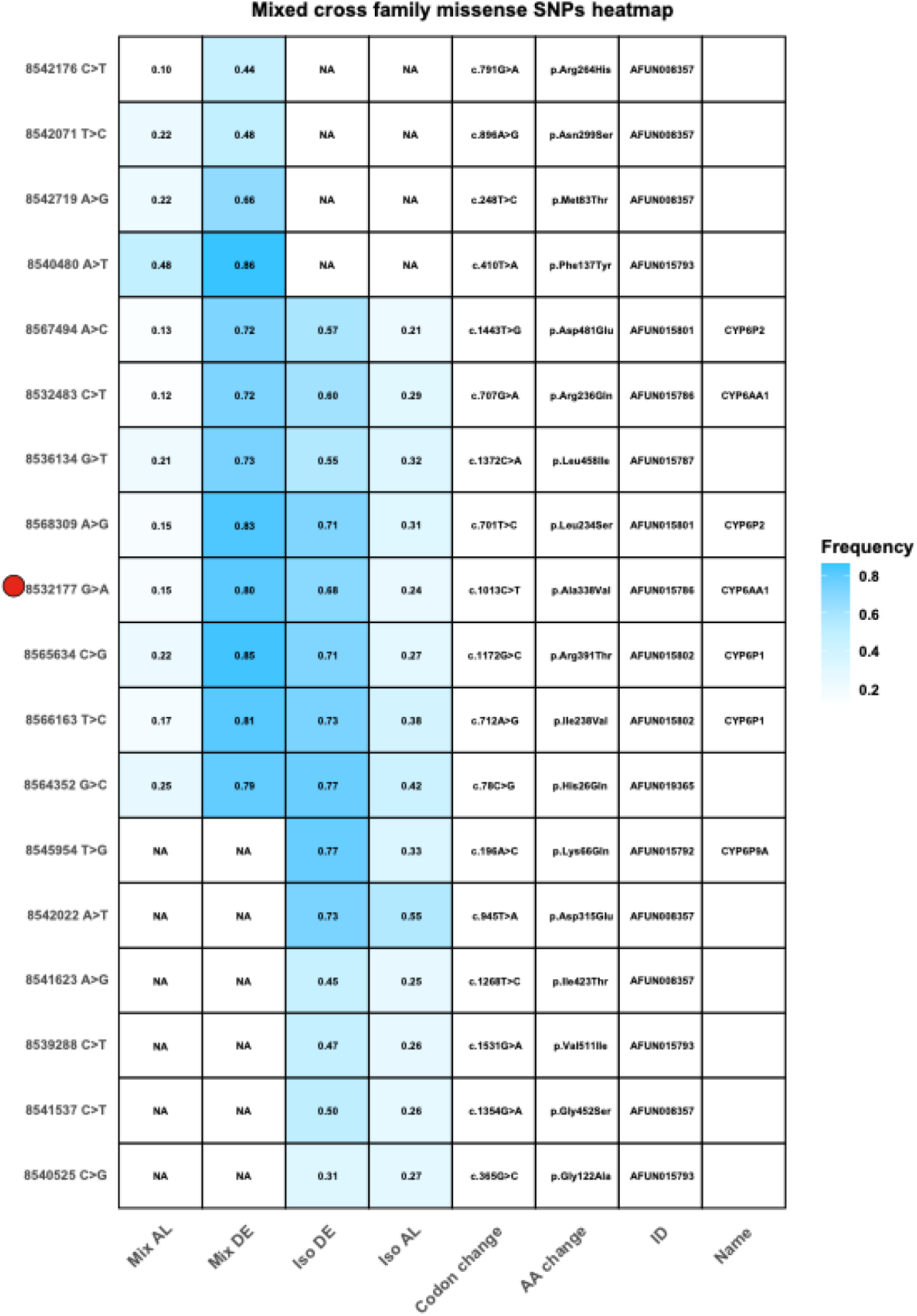
Heatmap of missense SNPs identified from the BSA in the F_7_ isofemale and mixed cross families in key detoxification genes in the *rap* QTL. All the missense SNPs identified in the *CYP6* cluster located on the negative strand also cause a missense mutation in the solute carrier family 8 (sodium/calcium exchanger), extending 130kb on the leading strand. Red dot highlights missense SNP in the *CYP6AA1* that was also identified in the QTL peak of conventional QTL mapping.

In addition to the polymorphism detected in the *CYP6* cluster, we have also detected polymorphism in ion channel genes within the *rap1* QTL, including sodium-calcium exchanger solute carrier family 8 (AFUN020405), sodium-potassium-transporting ATPase subunit alpha (AFUN008354), glutamate receptor ion channel gene (AFUN020159) and voltage-gated chloride channel gene (AFUN008007) (Dataset 8 and 9). Additionally, missense mutations were detected in carboxyl esterase genes AFUN015787 and AFUN015793 located within the *rap1* QTL (Figure S10). The role of these genes in pyrethroid resistance in *An. funestus* has not yet been explored.

Bulk segregant analyses (BSA) also revealed missense SNPs with significant allele frequency differences between the dead and alive groups in genes outside the *rap* QTL, including genes in the *CYP32* cluster, *CYP4* cluster and GSTU2 (AFUN008426) (Figure S10 and S11). Additionally, missense SNPs with significant allele frequency difference between dead and alive were detected in cuticular proteins (AFUN001523, AFUN001635, AFUN001203, AFUN021475, AFUN021476, AFUN004601). On the other hand, Indel-based BSA detected 317 indels with significant allele frequency differences within the *rap1* QTL in the F_7_ isofemale family and 293 in the F_7_ mixed-cross family.

Allele frequency differences for genes previously implicated in insecticide resistance, as well as genes potentially involved in resistance identified through BSA analyses, are presented as heatmaps in Figure S10 to S14. In this study, we highlight SNPs with significant allele frequency differences between susceptible and resistant populations, which can serve as molecular markers for surveillance of α-cypermethrin resistance.

## Discussion

Elucidating molecular mechanisms driving resistance to WHO-recommended insecticides is crucial for vector control and management strategies. This research highlights the discovery of a QTL conferring resistance to α-cypermethrin, named *rap1* QTL. Understanding the molecular basis of resistance to α-cypermethrin is crucial to determining the efficacy of ngITNs, especially those containing α-cypermethrin as an active ingredient.

In this study, we integrated two sequencing strategies to identify genomic regions in *An. funestus* associated with resistance to α-cypermethrin. Initially, to reduce the cost of sequencing while achieving sufficient genome-wide genotyping of segregating individuals, we employed double digest restriction-site associated DNA sequencing (ddRADseq) of an isofemale family segregating population at the F_2_ and the F_7_ generation. Secondly, we exploited a strategy of pooling DNA from individuals with extreme phenotypes to perform a cost-effective, rapid, and efficient QTL mapping discovery. Next-generation sequencing coupled with bulk segregate analysis (NGS-BSA) was performed on pools of Dead and Alive, segregating individuals from the F_7_ iso-female family and a mixed cross family independently. In our study, traditional QTL mapping provided a higher resolution for QTL mapping compared to BSA. Traditional QTL mapping focuses on recombination events between defined markers in a segregating population, identifying smaller and tightly linked QTLs. While BSA proved to be an efficient and rapid method for detecting genomic areas linked to resistance, however, with a lower resolution compared to traditional QTL mapping.

Traditional QTL mapping using F_7_ segregating individuals (F_7_ family) identified a 1.44 Mb QTL conferring resistance to α-cypermethrin in the *An. funestus*. Conversely, QTL mapping in the F_2_ segregating population (F_2_ family) did not detect a candidate region above the genome-wide threshold. This is due to increased recombination events by successive generations of sibling-mating in the F_7_ compared to the F_2_ family. This increase in recombination events is evident in the length of the genetic linkage map in the F_7_ family compared to the F_2_ Family (Figure 1). The *rap1* QTL when fit into a regression model explains only 22.5% when of the variance in susceptibility to α-cypermethrin. However, investigating a marker genotype on the QTL peak revealed a significant and strong association between resistant genotypes and α-cypermethrin resistance. The significant association between genotypes and resistance at the peak might reflect a more direct influence compared to the overall contribution of the QTL. The difference observed in the susceptibility to α-cypermethrin cannot be explained fully by the broader *rap1* QTL. Our analysis indicates there are other factors influencing the resistant phenotype. These factors may include other genetic loci with smaller effects that weren’t detected in our analysis. In earlier studies mapping genetic *A. funestus* loci associated with resistance to permethrin using microsatellite markers successfully identified *rp1* QTL at the F2 generation, explaining significant portion of phenotypic variance in pyrethroid resistance. While microsatellites genotyping demonstrated high resolution in QTL mapping detection, ddRADseq offers a more comprehensive genome-wide perspective. The dense single nucleotide variants provided by ddRADseq enabled a thorough investigation of the recombination landscape at the F_2_ and the F_7_ generation and allowed the discovery of missense SNPs with the highest LOD scores in the QTL peak in detoxification genes particularly *CYP6AA1* and *CYP6P2*. The choice between these methods should consider factors such as the genetic architecture of the trait, available resources, and the desired resolution of mapping. While in our analysis, we used a dense marker map to map *rap1* QTL, the sample size limited the detection of other QTLs with smaller effects. The detection of minor QTLs is inherently influenced by the sample size of the mapping population [25]. The sample size in our analysis was adequate to detect a major QTL associated with resistance to α-cypermethrin. However, increasing the sample size may facilitate the detection of additional QTLs with smaller effects.

The identified *rap1* QTL overlaps with the *rp1* QTL that was previously mapped to confer resistance to permethrin in *An. funestus* [12]. This overlap highlights the similarity in the molecular basis of resistance to Type I (permethrin) and Type II (α-cypermethrin) pyrethroids in this species. The association of *rp1* locus to permethrin resistance, particularly the CYP6 genes cluster, was demonstrated in several studies. Transcriptomic profiling of the *rp1* locus in the resistant FUMOZ colony revealed a significantly high expression of the duplicated *CYP6P9a* and *CYP6P9b*, highlighting their predominant role in resistance [13, 18]. The Resistant alleles of *CYP6P9a/b* in *An. funestus* were found to reduce the efficacy of insecticide-treated bed nets, particularly pyrethroid-only nets [4, 5, 26]. Additionally, Africa-wide genomic surveillance of *An. funestus* of the CYP6 gene cluster identified a signature of a selective sweep in southern Africa, revealing evidence of a major population division between southern Africa and elsewhere [16]. The CYP6 cluster contains complex structure variation Africa-wide with a gene copy number polymorphism and a 6.5 kb insertion between *CYP6P9a* and b [13]. This 6.5 kb insertion has been demonstrated to enhance P450-mediated resistance, where mosquitoes with the homozygote-resistant copy of the insertion have a greater chance of surviving exposure to pyrethroid-treated nets [8]. In our study, we found a strong correlation between resistant alleles of the 6.5 kb insertion and *CYP6P9a* gene with the chance to survive exposure to α-cypermethrin. These findings further indicate that the genetic basis underlying different classes of pyrethroids insecticide is broadly the same, and DNA-based assays developed to track resistance to type I permethrin around the CYP6 cluster are reproducible for type II α-cypermethrin.

Genetic linkage maps illustrate the difference in the recombination landscape between the F_7_ and the F_2_ families. The Marey plots illustrate the structural difference between the FANG and the FUMOZ genomes impacted by different successive generations of inbreeding and the geographical distance of the FANG and the FUMOZ strains (Figures 1 C and D). The high recombination rate (cM/Mb) in the F_7_ family is correlated with an increase in the number of cross-over events compared to the F_2_ family (Figure 2) (Table 1). In both families, recombination events were observed to occur infrequently near centromeric regions and were more common at the telomeric ends of chromosomes. This pattern is consistent with the tendency for recombination rates to be lower in centromeric or heterochromatic regions [27, 28]. However, due to the lower sequencing coverage near the centromeres in this study, recombination in these areas remains uncertain *in An. funestus*. Additionally, we did not specifically investigate whether recombination hotspots align with gene-dense regions, which would offer more insight into the evolutionary and functional significance of these regions in *An. funestus*. Genetic recombination in gene-dense regions accelerates genetic diversity by shuffling alleles, leading to a higher potential for adaptation to changing environments or selective pressures, such as insecticide resistance [21, 29].

Due to the advancement in sequencing technologies, well-assembled reference genomes reduced the necessity for traditional linkage maps. Linkage maps rely on recombination frequencies to determine the genetic distance of markers, therefore, when physical maps of genomes were not available or incomplete. In the era of high-quality reference genome assemblies, a linkage map is often redundant for QTL discovery. Identifying the molecular basis of a phenotype can be directly mapped into the genome by comparing allele frequencies between phenotypic groups, providing a precise mapping of candidate regions in the physical genome. Bulk segregant analysis (BSA) leverages the physical distance from the assembled genome to identify candidate regions by comparing allele frequencies between genomic pools of two extreme phenotypes. In our study, BSA provided a faster, cheaper and more straightforward approach with fewer computation steps to identify candidate regions while identifying SNPs in the candidate region that may have a deleterious effect on gene function. The performance of the BSA analysis in detecting candidate regions was assessed in the same F_7_ isofemale family used in traditional QTL mapping and an F_7_ mixed-cross family. The BSA approach on the F_7_ isofemale family detected a candidate region overlapping the *rap1* QTL, further supporting its role in α-cypermethrin resistance. The detected candidate region represents 6.9% of the total genome (14.36 Mb) compared to the *rap1* QTL ∼ 0.5% of the total genome size (1.44 Mb). While the detected candidate region is broad, the strongest peak overlaps with the *rap1* QTL. On the other hand, BSA analysis performed on the F_7_ mixed-cross family discovered many candidate regions, accounting for 31.1% of the total FUMOZ genome. The number of candidate regions identified in the mixed-cross family underscores the substantial genomic variation present in mixed populations. Nevertheless, the peak of the candidate region with the strongest signal overlaps with the *rap1* QTL, indicating the roles of the *rap1* QTL loci in resistance to α-cypermethrin in the overall resistant population, not specifically in the isofemale family. It is often challenging to generate isofemale families from field populations when crossed with the susceptible colonies to map resistant loci in field populations.

Current methods often use genome-wide association of pooled individuals (GWAS-PoolSeq) to identify resistant loci in field-resistant populations compared to susceptible populations from the same geography. However, this method is limited where resistance is high, reducing the ability to distinguish between resistant and field-susceptible populations [17]. To address these challenges, we propose generating a segregating population through a mixed cross between the field-resistant population and the laboratory-susceptible colony. Using BSA coupled with next-generation sequencing will provide a rapid, efficient and cost-effective approach to identify resistant loci in field resistant populations. BSA alone was not sufficient to pinpoint resistant markers since the discovered candidate regions were broad. To refine candidate regions, we propose integrating the DNA-based BSA analysis with RNA sequencing (RNAseq) to identify highly expressed genes within candidate regions. Alternatively, an RNA-based approach could be employed, allowing simultaneous identification of candidate regions and differentially expressed genes from the same RNA extraction. Furthermore, we recommend increasing the recombination events by successive generations of crossing, F_7_, in our study, generating large sample sizes for each pool, specifically selecting individuals with the most resistant and most susceptible phenotypes, as defined by exposure duration. These suggested strategies will enhance the power to identify key resistant markers rapidly, in a cost-efficient manner and with less effort, especially where resistance is rapidly developing and segregating susceptibility is no longer prevalent.

By utilising a well-characterized insecticide like α-cypermethrin, our study establishes a framework for future studies aimed at mapping resistance where resistance to new insecticide formulations evolves rapidly. This approach not only ensures a sustainable and adaptable strategy for resistance monitoring but also underscores the critical role of Bulk segregant analysis (BSA) as a tool for ongoing surveillance. Integrating BSA with traditional QTL mapping in our study identified polymorphism within the *rap1* QTL, primarily in key CYP450 genes (Table 3). While traditional QTL mapping identifies smaller and tightly linked QTLs, it does not provide a comprehensive view of polymorphism in that region because of the genome-reduced representation approach. BSA, on the other hand, highlights the polymorphism in candidate regions to accurately identify resistant markers that can be used in DNA-based susceptibility assay. In our study, we identified SNPs that are highly represented in the resistant segregating population compared to susceptible and vice versa that can be a target for susceptibility DNA-based assay (Table 3).

Further, BSA highlighted indels polymorphism within the *rap1*QTL that may have an impact on the resistant phenotype. Our results improve our understanding of the molecular basis of resistance to α-cypermethrin, inform the design of diagnostic tools and propose a cost-efficient surveillance approach to track and detect resistance across Africa.

## Materials and methods

### Mosquitoes rearing and crossing

Reciprocal crosses were established between *An. funestus* laboratory multi-resistant colony (FUMOZ) originating from Southern Mozambique and the insecticide susceptible colony (FANG) derived from Angola [30]. Crosses were performed in mass using a 1:1 ratio, with a minimum of 100 males from FANG and 100 females from FUMOZ. A reciprocal cross was also conducted, involving 100 females from FANG and 100 males from FUMOZ. Mated and blood-fed F_0_ females were individually allowed to lay F_1_ eggs using a forced oviposition technique to generate iso-female family lines [20, 21]. Insecticide bioassays were conducted using F_2_ generation of the iso-female lines. Females aged 2-5 days old were exposed to α-cypermethrin 1X (0.05%). Since there were no observable quantitative trait loci (QTL) at this generation, the progeny of each iso-female line was reared until F_7_ generation.

In a separate crossing experiment, F_1_ females were permitted to collectively lay eggs to create mixed cross families from the two reciprocal crosses between FANG and FUMOZ. As mentioned above, iso-female and mixed cross lines were reared till the F_7_ generation under controlled insectary conditions [12, 14, 22]. Insecticide bioassays were performed at the F_7_ generation following the standard WHO protocol. In brief, mosquitoes were exposed to 1X (0.05%) α-cypermethrin in WHO exposure tubes at two durations: 90 minutes (LT80) and 20 minutes (LT20).

Following exposure, mosquitoes were transferred to clean holding tubes, and mortality was recorded 24 hours post-exposure [3, 22–24]. Individuals that survived insecticide exposure after 90 minutes were categorized as highly resistant (HR), while those that died after 20 minutes of insecticide exposure were classified as highly susceptible (HS). Parental female founders of the iso-female families, along with progeny subjected to bioassays at the F2 and F7 generations, were collected and preserved in 80% ethanol for DNA extraction. To lyse mosquito tissues for DNA extraction, an individual mosquito was placed in each well of a 96-well plate with a180 ul of lysis buffer, 20 ul of proteinase K, and a metal ball bearing. Mosquitoes’ tissues were ground using a Tissue Lyser for 5 minutes at maximum frequency and then incubated in a preheated incubator at 56°C for 3 hours. following incubation plates were centrifuged at ∼2000 rpm in a plate centrifuge to collect condensation and liquid at the bottom of the tube. DNA extraction was conducted in 96 well-plates using DNeasy kit (Qiagen) according to manufacturer’s recommendation, eluted in 100 ul AE buffer and stored at -20° C. DNA quality was assessed using gel electrophoresis and concentration was measured using Quant-iT PicoGreen dsDNA Assay kit (Thermo Fisher, MA USA).

## DNA extraction and sequencing

### Double digest restriction-site associated DNA sequencing (ddRADseq)

Female progeny phenotyped according to the bioassay test from two isofemale families at the F_2_ and F_7_ generation originating from a FUMOZ female crossed with a FANG male were selected for ddRADseq. For each family, 96 samples were sequenced including 94 females (48 resistant and 46 susceptible) and the parental (F_0_) FANG and FUMOZ females. ddRADseq libraries were prepared by IGATech (Udine, Italy) using a custom protocol, with minor modifications to Peterson’s double digest restriction-site associated DNA preparation [31]. Genomic DNA was fluorometrically quantified, normalized to a uniform concentration and 75ng are double digested with 2.4U of both NlaIII and MluCI endonucleases (New England BioLabs) in 30μL reaction supplemented with CutSmart Buffer and incubated at 37°C for 90’, then at 65°C for 20’. Fragmented DNA was purified with 1.5 volumes of AMPureXP beads (Agencourt) and subsequently ligated with 200U of T4 DNA ligase (New England BioLabs) to 2pmol of both overhang barcoded adapters for NlaIII and MluCI cut sites in 40μL reaction incubated at 23°C for 60 min and at 20°C for 60 min followed by 20 min at 65°C. Samples were pooled into multiplexed batches and bead purified as before. For each pool, targeted fragments distribution is collected on BluePippin instrument (Sage Science Inc.) setting the range of 400 - 520 bp using a 2% agarose cassette. Gel eluted fractions were amplified with indexed primers using Phusion High-Fidelity PCR Master Mix (New England BioLabs) in as final volume of 50μL and subjected to the following thermal protocol [95°C, 3’] - [95°C, 30’’ - 60°C, 30’’ - 72°C, 45’’] x 12 cycles - [72°C, 2’]. Products were then purified with 1 volume of AMPureXP beads. The resulting libraries were checked with both Qubit 2.0 Fluorometer (Invitrogen, Carlsbad, CA) and Bioanalyzer DNA assay (Agilent Technologies, Santa Clara, CA). Libraries were then sequenced with 150 cycles in paired-end mode on NovaSeq 6000 instrument following the manufacturer’s instructions (Illumina, San Diego, CA). Sequencing data for the F_2_ family is available in ENA under accession number PRJEB82690 and for the F_7_ family under PRJEB82873.

### Pooled DNA sequencing (poolseq)

DNA pools from the F_7_ iso-female family (pooled 36 Dead and 48 Alive separately) and the F_7_ mixed cross family (pooled 96 Dead and 96 Alive separately) were constructed by pooling an equal amount of DNA from individual mosquitoes to produce a dead DNA pool and an alive DNA pool for each family. DNA pools were sequenced by Novogene (Cambridge, United Kingdom). Libraries were prepared using standard protocols to an average fragment size of 350 bp. Whole genome sequencing was performed using the NovaSeq 6000 Illumina platform with paired-end sequencing reads of 150 bp. The sequencing data is available in ENA under accession PRJEB70297.

### Assessment of correlation between CYP6P9a and 6.5 kb structural variant and α-cypermethrin resistance using molecular markers 6.5 kb intergenic structural variant genotyping

Individuals (alive and dead from α-cypermethrin bioassays) from the F_7_ isofemale family were individually genotyped for the 6.5 kb structural variant frequency to understand its association resistance to α-cypermethrin using the protocol described (Mugenzi et al., 2020). Genotyping is performed using a PCR assay which applies 3 primers, two of which flank the insertion region (FG_5F: CTTCACGTCAAAGTCCGTAT and FG_3R: TTTCGGAAAACATCCTCAA) a third which flanks the 6.5 kb region (FZ_INS5R: ATATGCCACGAAGGAAGCAG). Individuals harbouring the 6.5 kb structural variant have a band of 569 bp, whereas those without it have a band at 266 bp. Heterozygote individuals have two bands at both 569 bp and 266 bp after the Kapa Taq (Kapa Biosystems, Boston, MA, USA) PCR. A correlation analysis was then carried out to understand the association between survival to diagnostics doses of α-cypermethrin and the resistant allele frequency.

### Determination of *CYP6P9a* genotype associated with pyrethroid resistance

A previously discovered mutation in *CYP6P9a* implicated in pyrethroid resistance was assayed using fragment length polymorphism (RFLP) [4, 5]. Genotyping the frequency of the *CYP6P9a*_R alleles in the F_7_ isofemale family (96 individuals) was described previously (Weedall et al., 2019). Using a pair of forward (5’-TCCCGAAATACAGCCTTTCAG-3) and reverse (5’-ATTGGTGCCATCGCTAGAAG-3’) primers, partial *CYP6P9a* was amplified to include the region harbouring the A/G mutation in a 450 bp fragment using Kapa Taq PCR kits (Kapa Biosystems, Boston, MA, USA). The amplified fragment was digested (in a 2 h reaction at 65 °C) using Taq1α restriction enzyme that cuts the mutation sites yielding fragments of 350 bp and 100 bp for the resistant alleles while the susceptible alleles remain uncut. Odds ratios were calculated to establish the association between survival to α-cypermethrin and the frequency of the resistant allele (*CYP6P9a*-R allele) in the following manner [4, 5, 8, 19].

### Variant calling of ddRadseq

Pair-end illumina ddRADseq reads were demultiplexed using the process_redtags utility included in stacks (F_2_ v2.53, F_7_ v2.61)[32, 33]. A total of 468,893,296 and 1,030,538,210 reads were demultiplexed for the 96 individuals from the F_2_ and the F_7_ isofemale families respectively (Table S1). The quality of the demultiplexed reads was assessed with FastQC (v0.11.9) [34]. Reads were aligned to the *Anopheles funestus* AfunF3 reference genome (GeneBank assembly GCA_003951495.1) using bwa-mem2 (v 2.2.1) with default parameters and alignment sam files were converted to bam using samtools view (v 1.13) [35–39]. Sequenced loci from aligned reads were detected using the gstacks program included in Stacks (F_2_ v2.53, F_7_ v2.64) [32]. Variants in the aligned files were detected using the population program included in Stacks (F_2_ v2.53, F_7_ v2.64). Finally, variants were filtered using vcftools (F_2_ v0.1.16, F_7_ v0.1.15) based on a minimum depth of 10 (--minDP 10), a maximum missing value of 0.75 (--max-missing 0.75), minimum genotype quality of 30 (-- minGQ 30), keeping only bi-allelic SNPs and removing all indels [40]. Duplicated SNPs were removed using using bcftools (v1.21) [38, 41]. After filtering, 55,520 out of a possible 135,985 sites were kept in the F_2_ isofemale family and 285,204 variants were kept out of 1,591,234 sites in the F_7_ isofemale family. To identify polymorphic and monomorphic regions in the F_2_ and the F_7_ iso-female segregating population VCF files were imputed using beagle (v 5.4). Polymorphic and monomorphic sites were visualised independently in a density plot (Table S1). The variant calling pipeline is available in https://github.com/alyazeeditalal/RAD_seq_lepmap3-pipeline.

### Construction of Genetic Linkage Map

Genetic linkage maps for the F_2_ and the F_7_ families were constructed independently using Lep-MAP3 [42] as an “F_2_” population. To produce a pedigree file for the Lep- MAP3 pipeline, the F0 founders of the inter-cross were used as grandparents to assign two dummy F1 parents and the 64 segregating individuals were designated as F_2_ offspring, irrespective of the family generation of inter-crosses. The segregation ratio of markers is expected to reflect an F_2_ population (1:2:1), and the number of successive generations was implemented to increase recombination breakpoints. Parental markers were called from the VCF and pedigree files using the ParentCall2 module. Called markers were filtered using the Filtering2 module with a parameter to remove missing markers that are not present in at least 80% of the individuals (missingLimit=0.8) and remove markers with distorted segregation (dataTolerance = 0.0001).

Identified markers were assigned to linkage groups using the module SeperateChromosomes2 iteratively using a LOD score of 5-10. LOD score limit of 7 for the F_7_ family and LOD score limit of 5 for the F_2_ family (lodLimit) produced the best order. Parameters including removing linkage groups with a smaller number of markers than 100 (sizeLimit=100) and (lod3Mode= 2) were also used with the SeperateChromosomes2 module to split markers to the three major linkage groups. For the F_7_ family, 20,016 markers were separated into 3 major linkage groups whereas 18,570 markers were split into 3 major linkage groups in the F_2_ family. The module JoinSingle2All was called by testing several LOD limits, but no single markers were further added to the main 3 linkage groups in both families. Markers were ordered in each linkage group separately using the OrderMarkers2 module with the parameter grandparentPhase=1 to phase data according to (homozygote) grandparents and marker intervals were calculated (calculateIntervals = file). Marker ordering for each linkage group was run 5 times and the order with the highest likelihood was retained for each linkage group. Finally, phased output data from OrderMarker2 were converted to genotypes using map2genotypes.awk script. The marker dataset was manually filtered by aligning them with the corresponding chromosome in the FUMOZ reference genome. Specifically, markers in linkage group 1 were matched to chromosome AfunF3_2, markers in linkage group 2 to chromosome AfunF3_3, and those in linkage group 3 to chromosome AfunF3_X. This process resulted in 16,655 markers for the F_2_ family and 18,613 for the F_7_ family organised into three linkage groups corresponding to chromosomes (Table 1). The linkage map construction pipeline using Lep-MAP3 is available in https://github.com/alyazeeditalal/RAD_seq_lepmap3-pipeline. Then, using the (jittermap) function from the rqtl package, linkage maps were estimated to avoid placing markers at the same position [43]. The estimate of pairwise recombination fraction between markers and LOD score reflects the strength of linkage between each pair. A high LOD score, and a low recombination fraction indicate a strong linkage between each pair of markers, suggesting that they are physically close on a chromosome and thus more likely to be inherited together (Supplementary S2). The Asmap R package was used to visualise markers ratio and segregation distortion [44].

### QTL analysis

QTL mapping was conducted using rqtl using the final number of markers used to construct the genetic linkage map for both F_2_ and the F_7_ families separately [43]. Genetic linkage maps were investigated ensuring that markers with missing values in any of the individuals were removed and markers with high segregation distortions were removed according to the lep-MAP3 Filtering2 module where markers with an odd ratio of 1:10,000 (or a P-value threshold equivalent to 0.0001) were filtered out. The interval mapping genotyping probability was calculated using the kosambi map function with a 10 cM step size and an assumed error probability of 0.0001. The QTL was identified using a whole genome scan on all individuals with a single QTL model using Haley-Knott regression with a binary model where resistant mosquitoes were recorded as 1 and susceptible mosquitoes were recorded as 0. The result from the first scan was permuted 1000 times to obtain a genome-wide LOD score significant threshold at p-values < 0.05 corresponding to a LOD score of 3.63 for the F_2_ family and 3.82 for the F_7_ family. The QTL peak was identified by the function find.marker () provided by the rqtl package and then the QTL was fit in a model using Haley-Knott regression to estimate percent variance explained by the QTL [43].

### Pooled-template sequencing variant calling

Pair-end trimmed Pooled DNA sequences were examined for, contamination, low-quality reads and the presence of adapter sequences using Fastqc [45]. Reads in fastq files format were aligned against the AfunF3 *An. funestus* reference genome (GeneBank assembly GCA_003951495.1) using bwa-mem2 (v 2.2.1) and the sam alignment files were converted to binary alignment bam files using samtools (v 1.13) [36–39]. The quality of the bam alignment files was assessed using samtools flagstat and qualimap [36, 38]. Before variant calling, alignment bam files were pre- processed as follows: 1) files were sorted by position using samtools. 2) Duplicated reads were tagged using the genomic analysis toolkit (gatk v4.6) https://github.com/broadinstitute/gatk ‘MarkDuplicates’ and removed by setting the REMOVE_DUPLICATES option to true. 3) Read groups in the alignment bam were changed to match the sample ID using gatk AddOrReplaceReadGroups [46]. Variant calling was conducted using freebayes-parallel, to reduce the run time the number of alleles to consider was set to 4 using the option (use-best-alleles 4). The resulting variant calling files were annotated using snpeff [47] and subsequently filtered using bcftools by excluding variants with quality less than or equal to 20 (QUAL <=20), GT field is missing (FMT/GT=”./.”), genotype quality less than 99 (GQ<99), read depth less than 100 (FMT/DP<100), keeping only biallelic variants and producing separate vcf files for SNPs and indels [38]. Finally, variants were converted to a table using gatk (v4.6) VariantsToTable to be used in the bulk segregant analysis (BSA).

### Bulk segregate analysis (BSA) and population genetics

BSA analysis was conducted using pools of dead and alive from the F_7_ isofemale family and the F_7_ mixed cross family. A custom bash script based on the QTLseqr R package was developed to run the BSA analysis [22]. BSA custom script and the complete poolseq pipeline is available here https://github.com/alyazeeditalal/Poolseq_analysis. SNPs were filtered before running the BSA analysis based on reference allele frequency of 0.20, minimum total depth equal to 100, maximum total depth equal to 400 and minimum sample depth of 40. The difference in allele frequency between alive and dead bulk (delta SNP-index) was calculated on a sliding window size of 1 Mb, while calculating the number of SNPs per window [23]. Confidence intervals at 95% and 99% were estimated using the quantile from simulating the delta-SNP-index per bulk over 1000 replications. Additionally, G-statistics was calculated genome-wide and a tricube-smoothed G-statistics (G’), a smoothing method on the G value across the genome, was calculated in a sliding window of 1 Mb [24]. G-statistics measure the goodness of fit between observed and expected values, assessing whether deviations are due to a chance or associated with a phenotype of interest, and determining if the observed differences are random or biologically relevant. A p-value was determined by comparing the G’ values to a non-parametrically estimated null distribution, with the assumption that the distribution of the G’ is a proximately log-normal. Negative log10 p-value and Benjamini-Hochberg adjusted p-values were calculated [48]. Candidate regions were identified using a genome-wide false discovery rate (FDR) of 0.01. Population differentiation (Fst) between dead and alive pools was calculated at the SNP level, a sliding window size of 1 Mb and gene-wise using grenedalf (https://github.com/lczech/grenedalf) [49].

## References

1. Organization, W.H., World malaria report 2023. 2023: World Health Organization.

2. Bhatt, S., et al., The effect of malaria control on Plasmodium falciparum in Africa between 2000 and 2015. Nature, 2015. 526(7572): p. 207-211.

3. Riveron, J.M., et al., Rise of multiple insecticide resistance in Anopheles funestus in Malawi: a major concern for malaria vector control. Malar J, 2015. 14: p. 344.

4. Weedall, G.D., et al., A cytochrome P450 allele confers pyrethroid resistance on a major African malaria vector, reducing insecticide-treated bednet efficacy. Sci Transl Med, 2019. 11(484).

5. Mugenzi, L.M.J., et al., Cis-regulatory CYP6P9b P450 variants associated with loss of insecticide-treated bed net efficacy against Anopheles funestus. Nat Commun, 2019. 10(1): p. 4652.

6. Hemingway, J., et al., Averting a malaria disaster: will insecticide resistance derail malaria control? Lancet, 2016. 387(10029): p. 1785–8.

7. Ibrahim, S.S., et al., Allelic Variation of Cytochrome P450s Drives Resistance to Bednet Insecticides in a Major Malaria Vector. PLOS Genetics, 2015. 11(10): p. e1005618.

8. Mugenzi, L.M.J., et al., A 6.5-kb intergenic structural variation enhances P450-mediated resistance to pyrethroids in malaria vectors lowering bed net efficacy. Molecular Ecology, 2020. 29(22): p. 4395–4411.

9. Hemingway, J., The way forward for vector control. Science, 2017. 358(6366): p. 998-999.

10. Böhmert, A.L., et al., A descriptive review of next-generation insecticide-treated bed nets for malaria control. Frontiers in Malaria, 2024. 2.

11. Dhadialla, T.S., G.R. Carlson, and D.P. Le, New insecticides with ecdysteroidal and juvenile hormone activity. Annu Rev Entomol, 1998. 43: p. 545–69.

12. Wondji, C.S., et al., Mapping a Quantitative Trait Locus (QTL) conferring pyrethroid resistance in the African malaria vector Anopheles funestus. BMC Genomics, 2007. 8(1): p. 34.

13. Wondji, C.S., et al., RNAseq-based gene expression profiling of the Anopheles funestus pyrethroid-resistant strain FUMOZ highlights the predominant role of the duplicated CYP6P9a/b cytochrome P450s. G3 Genes|Genomes|Genetics, 2021. 12(1).

14. Wondji, C.S., et al., Two duplicated P450 genes are associated with pyrethroid resistance in Anopheles funestus, a major malaria vector. Genome Res, 2009. 19(3): p. 452–9.

15. Barnes, K.G., et al., Genomic Footprints of Selective Sweeps from Metabolic Resistance to Pyrethroids in African Malaria Vectors Are Driven by Scale up of Insecticide-Based Vector Control. PLOS Genetics, 2017. 13(2): p. e1006539.

16. Weedall, G.D., et al., An Africa-wide genomic evolution of insecticide resistance in the malaria vector Anopheles funestus involves selective sweeps, copy number variations, gene conversion and transposons. PLOS Genetics, 2020. 16(6): p. e1008822.

17. Hearn, J., et al., Multi-omics analysis identifies a CYP9K1 haplotype conferring pyrethroid resistance in the malaria vector Anopheles funestus in East Africa. Mol Ecol, 2022. 31(13): p. 3642–3657.

18. Al-Yazeedi, T., et al., Overexpression and nonsynonymous mutations of UDP-glycosyltransferases are potentially associated with pyrethroid resistance in Anopheles funestus. Genomics, 2024. 116(2): p. 110798.

19. Mugenzi, L.M.J., et al., The duplicated P450s CYP6P9a/b drive carbamates and pyrethroids cross-resistance in the major African malaria vector Anopheles funestus. PLoS Genet, 2023. 19(3): p. e1010678.

20. Kurlovs, A.H., et al., Trait mapping in diverse arthropods by bulked segregant analysis. Current Opinion in Insect Science, 2019. 36: p. 57–65.

21. Al-Yazeedi, T., et al., The contribution of an X chromosome QTL to non-Mendelian inheritance and unequal chromosomal segregation in Auanema freiburgense. Genetics, 2024. 227(1).

22. Mansfeld, B.N. and R. Grumet, QTLseqr: An R Package for Bulk Segregant Analysis with Next-Generation Sequencing. The Plant Genome, 2018. 11.

23. Takagi, H., et al., QTL-seq: rapid mapping of quantitative trait loci in rice by whole genome resequencing of DNA from two bulked populations. Plant J, 2013. 74(1): p. 174–83.

24. Magwene, P.M., J.H. Willis, and J.K. Kelly, The Statistics of Bulk Segregant Analysis Using Next Generation Sequencing. PLOS Computational Biology, 2011. 7(11): p. e1002255.

25. Wang, H., et al., Effect of population size and unbalanced data sets on QTL detection using genome-wide association mapping in barley breeding germplasm. Theor Appl Genet, 2012. 124(1): p. 111–24.

26. Menze, B.D., et al., Experimental Hut Trials Reveal That CYP6P9a/b P450 Alleles Are Reducing the Efficacy of Pyrethroid-Only Olyset Net against the Malaria Vector Anopheles funestus but PBO-Based Olyset Plus Net Remains Effective. Pathogens, 2022. 11(6): p. 638.

27. Juneja, P., et al., Assembly of the genome of the disease vector Aedes aegypti onto a genetic linkage map allows mapping of genes affecting disease transmission. PLoS Negl Trop Dis, 2014. 8(1): p. e2652.

28. Boyle, J.H., et al., A Linkage-Based Genome Assembly for the Mosquito Aedes albopictus and Identification of Chromosomal Regions Affecting Diapause. Insects, 2021. 12(2).

29. Genetic diversity of the African malaria vector Anopheles gambiae. Nature, 2017. 552(7683): p. 96-100.

30. Hunt, R.H., et al., Laboratory selection for and characteristics of pyrethroid resistance in the malaria vector Anopheles funestus. Med Vet Entomol, 2005. 19(3): p. 271–5.

31. Peterson, B.K., et al., Double Digest RADseq: An Inexpensive Method for De Novo SNP Discovery and Genotyping in Model and Non-Model Species. PLOS ONE, 2012. 7(5): p. e37135.

32. Rochette, N.C., A.G. Rivera-Colón, and J.M. Catchen, Stacks 2: Analytical methods for paired-end sequencing improve RADseq-based population genomics. Mol Ecol, 2019. 28(21): p. 4737–4754.

33. Rochette, N.C. and J.M. Catchen, Deriving genotypes from RAD-seq short-read data using Stacks. Nature Protocols, 2017. 12(12): p. 2640–2659.

34. Andrews, S. FastQC: a quality control tool for high throughput sequence data. 2010; Available from: http://www.bioinformatics.babraham.ac.uk/projects/fastqc.

35. Ghurye, J., et al., A chromosome-scale assembly of the major African malaria vector Anopheles funestus. Gigascience, 2019. 8(6).

36. Li, H. and R. Durbin, Fast and accurate short read alignment with Burrows-Wheeler transform. Bioinformatics, 2009. 25(14): p. 1754–60.

37. Vasimuddin, M., et al. Efficient Architecture-Aware Acceleration of BWA-MEM for Multicore Systems. in 2019 IEEE International Parallel and Distributed Processing Symposium (IPDPS). 2019.

38. Danecek, P., et al., Twelve years of SAMtools and BCFtools. GigaScience, 2021. 10(2).

39. Li, H., et al., The Sequence Alignment/Map format and SAMtools. Bioinformatics (Oxford, England), 2009. 25(16): p. 2078–2079.

40. Danecek, P., et al., The variant call format and VCFtools. Bioinformatics, 2011. 27(15): p. 2156–2158.

41. Narasimhan, V., et al., BCFtools/RoH: a hidden Markov model approach for detecting autozygosity from next-generation sequencing data. Bioinformatics (Oxford, England), 2016. 32(11): p. 1749–1751.

42. Rastas, P., Lep-MAP3: robust linkage mapping even for low-coverage whole genome sequencing data. Bioinformatics, 2017. 33(23): p. 3726–3732.

43. Broman, K.W., et al., R/qtl: QTL mapping in experimental crosses. Bioinformatics, 2003. 19(7): p. 889–90.

44. Taylor, J. and D. Butler, R Package ASMap: Efficient Genetic Linkage Map Construction and Diagnosis. Journal of Statistical Software, 2017. 79(6): p. 1–29.

45. Riva, A., et al., Streamlining DNA Sequencing and Bioinformatics Analysis Using Software Containers. J Biomol Tech, 2019. 30(Suppl): p. S38–s39.

46. McKenna, A., et al., The Genome Analysis Toolkit: a MapReduce framework for analyzing next-generation DNA sequencing data. Genome research, 2010. 20(9): p. 1297–1303.

47. Cingolani, P., et al., A program for annotating and predicting the effects of single nucleotide polymorphisms, SnpEff: SNPs in the genome of Drosophila melanogaster strain w1118; iso-2; iso-3. Fly (Austin), 2012. 6(2): p. 80-92.

48. Benjamini, Y. and Y. Hochberg, Controlling the False Discovery Rate: A Practical and Powerful Approach to Multiple Testing. Journal of the Royal Statistical Society: Series B (Methodological), 1995. 57(1): p. 289–300.

49. Czech, L., J. Spence, and M. Exposito-Alonso, grenedalf: population genetic statistics for the next generation of pool sequencing. 2023.

