## Supplemental Data for "Genetic mapping of resistance: A QTL and associated polymorphism conferring resistance to alpha-cypermethrin in *Anopheles funestus*"

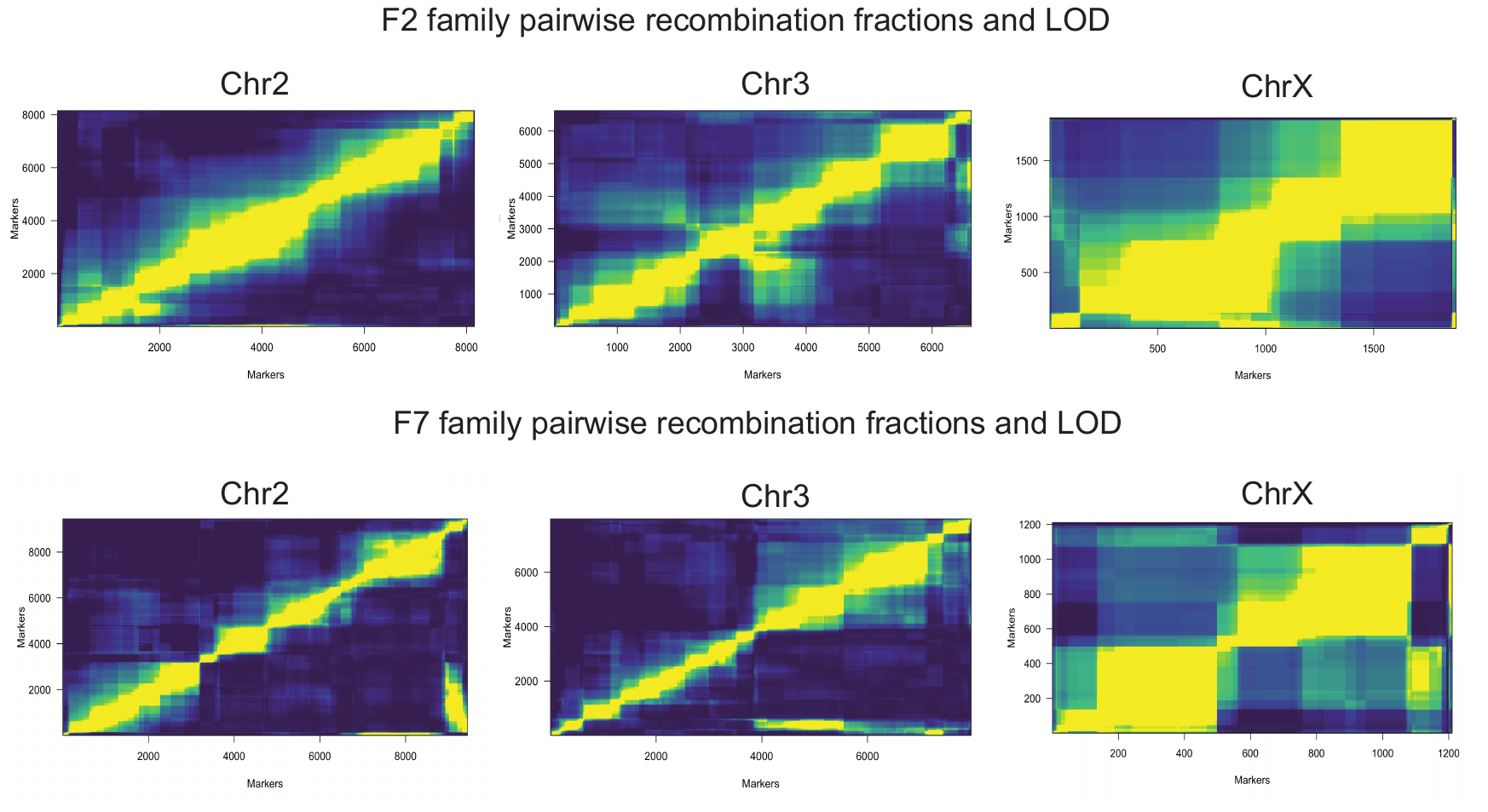

**Figure S1. Markers pairwise recombination fraction and LOD score per chromosome in the F2 and F7 isofemale family by Chromosome.** Plots of estimated recombination fractions (upper-left triangle) and LOD scores (lower-right triangle) for all pairs of markers in each chromosome, F2 isofemale family top half and F7 isofemale family bottom half. Yellow indicates linked (large LOD score or small recombination fraction) and blue indicates no linkage (small LOD score or large recombination fraction).

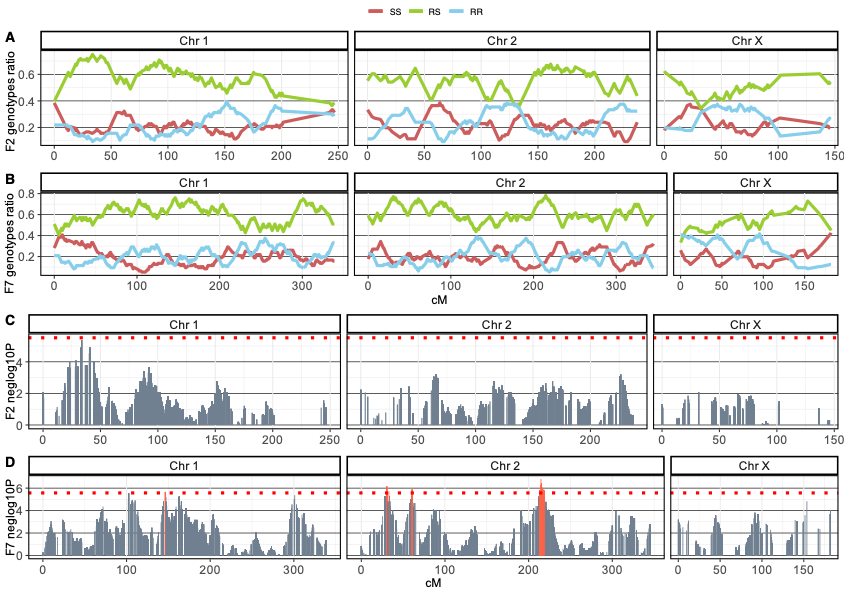

**Figure S2. Markers genotype ratio and segregation distortion in the F2 and the F7 family genetic linkage map.** The ratio of the SS genotype (Homozygote for FANG) in red, the RS genotype (heterozygote for FANG and FUMOZ) and the RR genotype (Homozygote for FUMOZ) for each marker in all individuals in the F2 (**A**) and F7 (**B**) isofemale families were plotted against the recombination distance (cM) of each family on the x-axis. Markers segregation distortion from the expected ratio (- log (10) p-value) in the F2 (**C**) and F7 (**D**) families plotted against the recombination distance (cM) of each family indicates that most markers segregation distortion is lower than the family-wide Bonferroni adjusted alpha level (red dotted line) of grey columns. While markers with a high segregation distortion are highlighted with red columns.

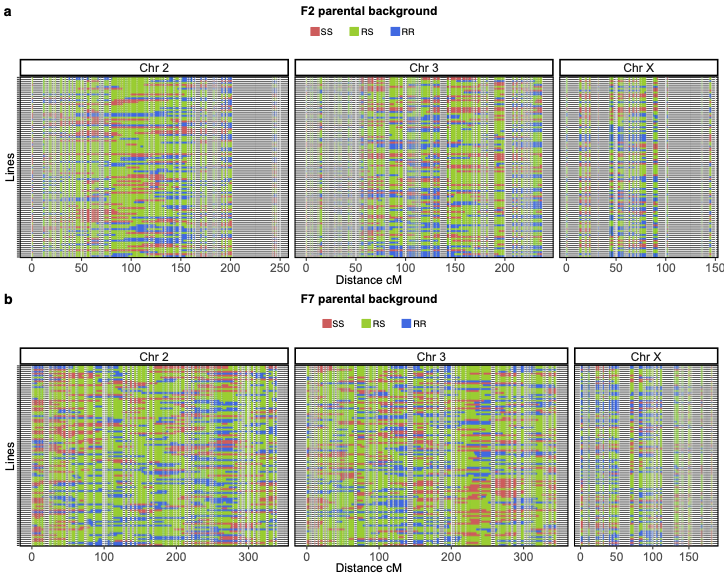

**Figure S3. Parental background of genotyped markers in all individuals in the F2 and the F7 isofemale families across the genetic distance.** Marker genotypes of segregating individuals (y-axis) were plotted against the genetic distance in cM (X-axis) in the F2 isofemale family (**a**) and the F7 isofemale family (**b**). Markers with the susceptible FANG background are coloured in red (SS), Markers with the resistant FUMOZ background are coloured in blue (RR) and markers that are heterozygote for both backgrounds (RS) are coloured in green

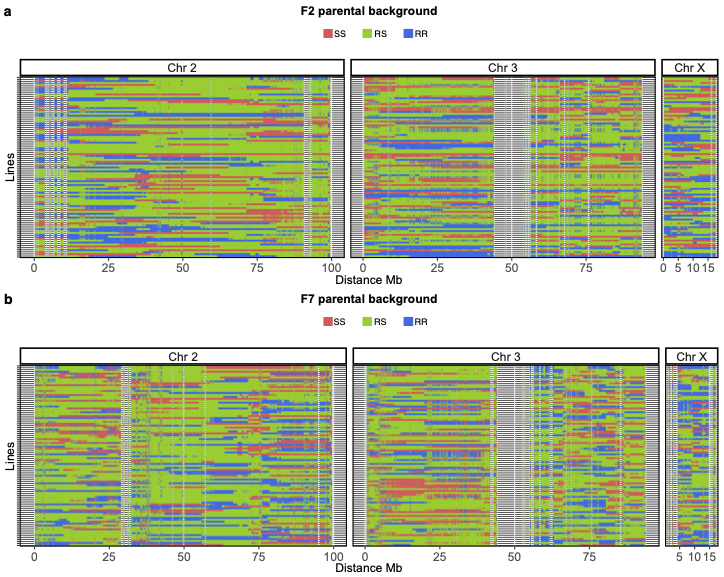

**Figure S4. Markers’ parental background in all individuals in the F2 and the F7 isofemale families across the physical distance.** Marker genotypes of segregating individuals (y-axis) were plotted against the FUMOZ reference physical distance in Mb (X-axis) in the F2 isofemale family (**a**) and the F7 isofemale family (**b**). Markers with the susceptible FANG background are coloured in red (SS), Markers with the resistant FUMOZ background are coloured in blue (RR), and markers that are heterozygote for both backgrounds (RS) are coloured in green.

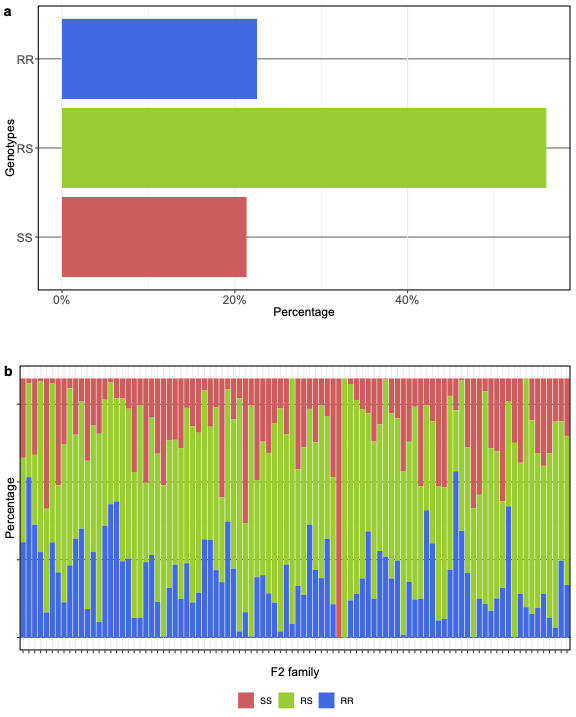

**Figure S5. Percentage of parental genotypes in the F2 family overall genetic linkage map and per the family individuals.** The plot highlights the percentages of marker genotypes in the overall genetic linkage map (a) and the percentage of genotypes per individual (x-axis) (b).

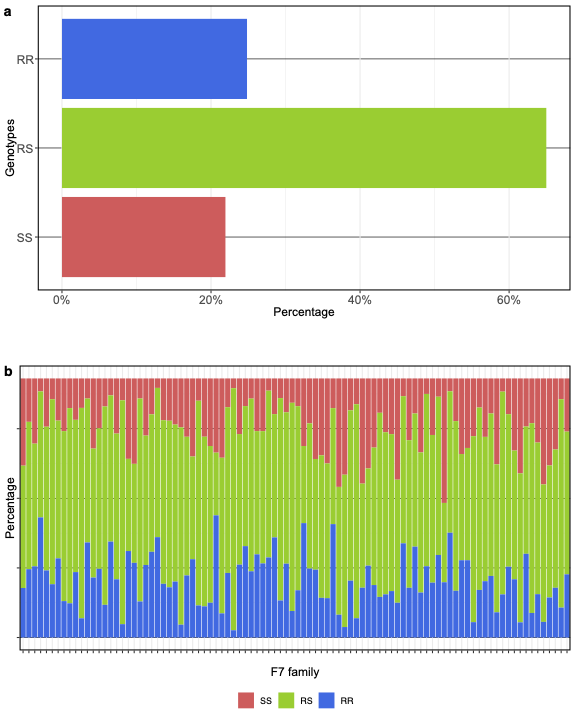

**Figure S6. Percentage of parental genotypes in the F7 family overall genetic linkage map and per the family individuals.** The plot highlights the percentages of marker genotypes in the overall genetic linkage map (a) and the percentage of genotypes per individual (x-axis) (b).

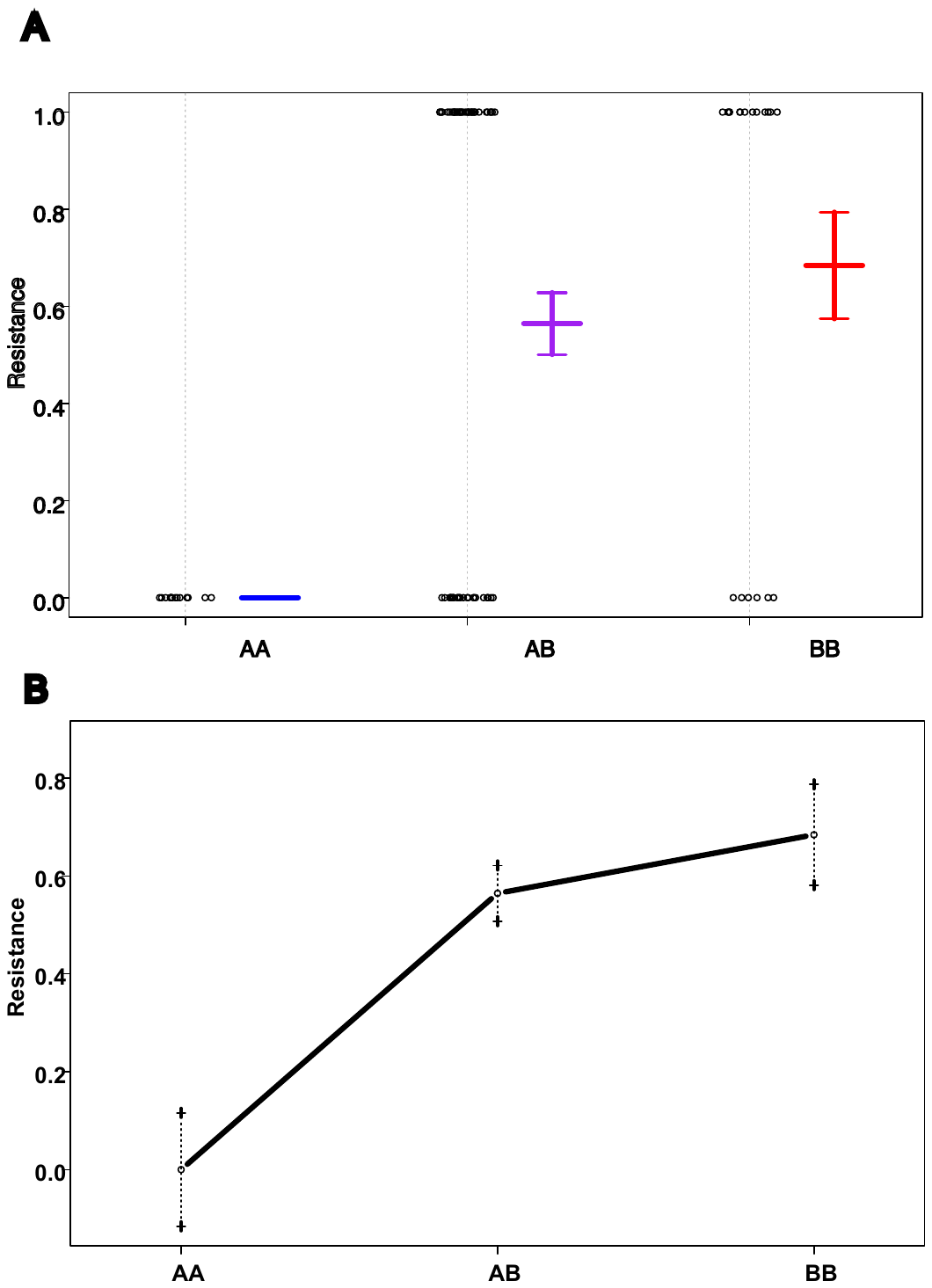

**Figure S7. The mean for the resistance phenotype.** The mean for the resistance phenotype in individuals with homozygous FANG genotype (AA) at the QTL peak is 0, compared to a mean of 0.68 for the resistant phenotype in individuals homozygote for FUMOZ genotype (BB), whereas the mean for the resistant phenotype in individuals with a heterozygote QTL peak is 0.56.

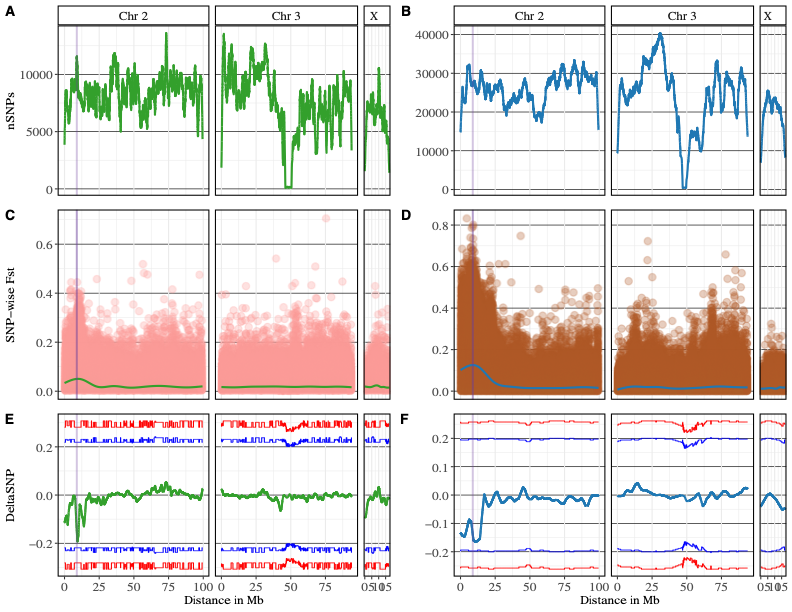

**Figure S8: SNP Density, genetic differentiation and delta SNP-index.** The number of SNPs (y-axis) in a window size of 1Mb was plotted genome-wide in the F7 isofemale family (A) and F7 mixed cross family (B). SNPs level F_st_ differentiation (y-axis) illustrates an elevated differentiation around the *rap* QTL locus (vertical line in purple) in the F7 isofemale family (C) and the F7 mixed family (D). Delta SNP-index was calculated in a window size of 1Mb (y-axis) in F7 isofemale family (E) and F7 mixed cross family (F). The 95 and 99 confidence intervals are represented by blue and red, respectively. The *rap* QTL locus is highlighted by the purple vertical line in all plots.

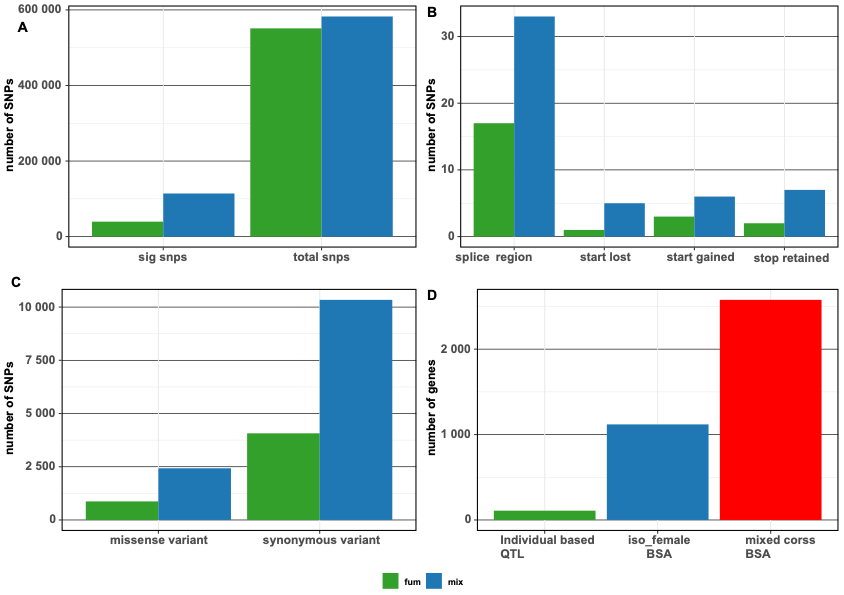

**Figure S9. The number of SNPs identified in BSA analysis.**

**Table S1. Summarised sequencing and variants statistics for ddRADseq**

| Family | Sequenced and genotyped samples | Total demultiplexed Reads | Number of SNPs called | Number of SNPs after filtering based on depth and quality | Polymorphic SNPs | Monomorphic SNPs | Final number of markers used in linkage map |
| --- | --- | --- | --- | --- | --- | --- | --- |
| F2 isofemale family | 96 | 468,893,296 | 135,985 | 55,520 | 46,160 | 9,360 | 16,655 |
| F7 isofemale family | 96 | 1,030,538,210 | 1,591,234 | 285,204 | 59,161 | 5,119 | 18,613 |

**Table S2. Genotype frequency in the segregating population from the F7 isofemale of the 6.5Kb insertion and CYP6p9a individually and together.**

| Marker | phenotype | RR% | RS% | SS% |
| --- | --- | --- | --- | --- |
| 6.5Kb SV | Dead | 15.8 | 45.6 | 95 |
|  | Alive | 84.2 | 54.4 | 5 |
| CYP6p9a | Dead | 1.5 | 47.4 | 94.7 |
|  | Alive | 85 | 52.6 | 5.3 |
| 6.5Kb SV/CYP6p9a | Dead | 15.8 | 44.6 | 95.2 |
|  | Alive | 84.2 | 55.4 | 4.8 |

**Table S3. Estimation of odd ratio for survivability amongst the F7 segregating progeny based on the inherited genotypes.**

|  |  | **RRvsSS** | **RRvsRS** | **RSvsSS** |
| --- | --- | --- | --- | --- |
| 6.5Kb SV | **Odd Ratio** | 101.33 | 4.4731 | 22.6538 |
|  | **Pvalue** | P=0.0001 | P = 0.0283 | P = 0.0032 |
|  | **Confidence interval** | 9.5784 to 1072.0396 | 1.1727 to 17.0621 | 2.8376 to 180.8557 |
| CYP6p9a | **Odd Ratio** | 102 | 5.1 | 20 |
|  | **Pvalue** | P = 0.0001 | P = 0.0166 | P = 0.0048 |
|  | **Confidence interval** | 9.6473 to 1078.4314 | 1.3448 to 19.3409 | 2.4992 to 160.0492 |
| 6.5Kb SV/CYP6p9a | **Odd Ratio** | 106.6667 | 4.3011 | 24.8 |
|  | **Pvalue** | P = 0.0001 | P = 0.0330 | P = 0.0024 |
|  | **Confidence interval** | 10.1042 to 1126.0461 | 1.1251 to 16.4421 | 3.1096 to 197.7894 |

**Table S4. BSA result using SNPs from F7 isofemale line.**

| Chr | Chr 2 |
| --- | --- |
| QTL | 1 |
| start | 183 |
| end | 14,572,176 |
| length | 14,571,993 |
| nSNPs in the resion | 11,8534 |
| Average SNPs Mb | 8,134 |
| Peak DeltaSNP | -0.19 |
| Pos Peak DeltaSNP | 9,326,280 |
| Avg DeltaSNP | -0.0781 |
| Max Gprime | 19.05 |
| Pos MaxGprime | 10,880,716 |
| Mean Gprime | 10.15 |
| Sd Gprime | 3.92 |
| Mean Pval | 1.75E-05 |
| Mean Qval | 0.00027 |

**Table S5. BSA results using SNPs for the F7 mixed cross line.**

| Chromosome | 2 | 3 | 3 | 3 |
| --- | --- | --- | --- | --- |
| QTL | 1 | 2 | 3 | 4 |
| start | 10,337 | 9,701,042 | 61,529,533 | 68,519,595 |
| end | 32,903,881 | 14,041,521 | 61,703,920 | 70,502,099 |
| length | 32,893,544 | 4,340,479 | 174,387 | 1,982,504 |
| nSNPs in the resion | 287,641 | 37,681 | 988 | 16,014 |
| Average SNPs Mb | 8,745 | 8,681 | 5,666 | 8,078 |
| Peak DeltaSNP | -0.1655684 | 0.04053316 | -0.039152 | -0.0075792 |
| Pos Peak DeltaSNP | 10,890,283 | 14,041,521 | 61,583,917 | 68,519,595 |
| Avg DeltaSNP | -0.0741001 | 0.02627342 | -0.0388078 | -3.20E-05 |
| Max Gprime | 45.1658105 | 3.93470615 | 3.61297417 | 3.8121426 |
| Pos MaxGprime | 7,781,510 | 11,730,785 | 61,583,917 | 68,913,760 |
| Mean Gprime | 20.6 | 3.8 | 3.6 | 3.7 |
| Sd Gprime | 11.7 | 0.1 | 0.013 | 0.06 |
| Mean Pval | 1.08E-05 | 0.0007071 | 0.00178292 | 0.00117832 |
| Mean Qval | 5.94E-05 | 0.00393495 | 0.00917292 | 0.00627456 |

**Table S6. Predicted effect of significant SNPs identified from BSA.**

| Effect | Mixed cross | Iso-female |
| --- | --- | --- |
| 3 prime UTR variant | 5619 | 1948 |
| 5 prime UTR premature start codon gain variant | 405 | 137 |
| 5 prime UTR variant | 2893 | 1138 |
| downstream gene variant | 43277 | 16857 |
| intergenic region | 39989 | 12830 |
| intragenic variant | 16 | 17 |
| Intron variant | 73710 | 26634 |
| missense variant | 2428 | 872 |
| missense variant & splice region variant | 15 | 7 |
| Splice acceptor variant & intron variant | 7 | 1 |
| Splice donor variant & intron variant | 1 | 0 |
| splice region variant | 33 | 17 |
| splice region variant & intron variant | 467 | 193 |
| splice region variant & stop retained variant | 61 | 0 |
| splice region variant & synonymous variant | 5 | 32 |
| Start lost | 6 | 1 |
| Stop gained | 2 | 3 |
| Stop lost | 7 | 0 |
| Stop lost & splice regionvariant | 5619 | 1 |
| Stop retained variant | 405 | 2 |
| synonymous variant | 10339 | 4070 |

**Table S7. BSA using indels for the F7 isofemale family.**

| CHROM | Chr 2 |
| --- | --- |
| qtl | 1 |
| start | 183 |
| end | 14,572,176 |
| length | 14,571,993 |
| nSNPs | 118534 |
| avgSNPs Mb | 8134 |
| peakDeltaSNP | -0.1932973 |
| posPeakDeltaSNP | 9326280 |
| avgDeltaSNP | -0.0781873 |
| maxGprime | 19.0527828 |
| posMaxGprime | 10880716 |
| meanGprime | 10.1500322 |
| sdGprime | 3.92494298 |
| meanPval | 1.75E-05 |
| meanQval | 0.00026652 |

**Table S8. BSA using indels for the F7 mixed-cross family**

| CHROM | Chr 2 | Chr 3 | Chr 3 | Chr 3 | Chr 3 | Chr X | Chr X |
| --- | --- | --- | --- | --- | --- | --- | --- |
| Candidate region | 1 | 2 | 3 | 4 | 5 | 6 | 7 |
| start | 11551 | 10861232 | 13300103 | 61561984 | 67955473 | 14788649 | 17497797 |
| end | 32526522 | 12319067 | 14665219 | 62757939 | 70594413 | 15210703 | 17655384 |
| length | 32514971 | 1457835 | 1365116 | 1195955 | 2638940 | 422054 | 157587 |
| nSNPs | 21586 | 912 | 951 | 540 | 1430 | 303 | 115 |
| avgSNPs Mb | 664 | 626 | 697 | 452 | 542 | 718 | 730 |
| peakDeltaSNP | -0.208 | 0.031 | 0.044 | -0.094 | 0.017 | -0.078 | -0.051 |
| posPeakDeltaSNP | 10113526 | 12319067 | 13300103 | 61591533 | 70594413 | 15210703 | 17655384 |
| avgDeltaSNP | -0.083 | 0.021 | 0.039 | -0.074 | 0.004 | -0.074 | -0.050 |
| maxGprime | 46.04 | 4.18 | 4.05 | 4.43 | 4.28 | 3.58 | 3.66 |
| posMaxGprime | 7006760 | 11736586 | 13933437 | 62310784 | 68873686 | 14897559 | 17655384 |
| meanGprime | 19.15 | 3.85 | 3.76 | 3.92 | 3.91 | 3.54 | 3.60 |
| sdGprime | 10.904 | 0.185 | 0.166 | 0.223 | 0.243 | 0.022 | 0.036 |
| meanPval | 4E-06 | 6E-04 | 9E-04 | 5E-04 | 6E-04 | 2E-03 | 1E-03 |
| meanQval | 2E-05 | 3E-03 | 5E-03 | 3E-03 | 3E-03 | 9E-03 | 7E-03 |

**Table S9. Indels number**

|  | F7 mixed cross Family | F7 Isofemale Family |
| --- | --- | --- |
| Total number of indels used in BSA | 44,482 | 41,676 |
| Total number of significant Indels | 8,613 | 3,146 |
| 3 prime UTR variant | 754 | 244 |
| 5 prime UTR variant | 289 | 125 |
| Conservative inframe deletion | 7 | 2 |
| Conservative inframe insertion | 21 | 9 |
| Disruptive inframe deletion | 19 | 1 |
| Disruptive inframe insertion | 10 | 4 |
| Downstream gene variant | 2,790 | 1,049 |
| Frameshift variant | 10 | 8 |
| Frameshift variant & splice region variant | 1 | 1 |
| Frameshift variant & start lost | 1 | NA |
| intergenic region | 3,052 | 1,066 |
| intron variant | 6,168 | 2,354 |
| noncoding transcript variant | 2 | NA |
| splice donor variant & intron variant | 1 | NA |
| splice donor variant & splice region variant & intron variant | 2 | NA |
| splice region variant | 6 | 1 |
| splice region variant & downstream gene variant | 1 | NA |
| splice region variant & intron variant | 35 | 11 |
| start retained variant | 1 |  |
| upstream gene variant | 2,845 | 1,121 |

**Table S10. Missense SNPs in the *rap1* QTL identified by BSA in the F7 isofemale family.** In the F7 isofemale family, we identified 14 missense SNPs in 9 different genes.

| **Effect** | **POS** | **REF** | **ALT** | **SNPindex.LOW** | **SNPindex.HIGH** | **deltaSNP** | **negLog10Pval** | **codon** | **Amino acid** | **ID** | **Name** | **description** |
| --- | --- | --- | --- | --- | --- | --- | --- | --- | --- | --- | --- | --- |
| Missense | 8532177 | G | A | 0.682 | 0.242 | -0.44 | 8.112 | c.1013C>T | p.Ala338Val | AFUN015786 | CYP6AA1 | Cytochrome P450 |
|  | 8532483 | C | T | 0.603 | 0.286 | -0.317 | 8.113 | c.707G>A | p.Arg236Gln | AFUN015786 | CYP6AA1 | Cytochrome P450 |
|  | 8536134 | G | T | 0.545 | 0.319 | -0.226 | 8.119 | c.1372C>A | p.Leu458Ile | AFUN015787 | NA | Carboxylic ester hydrolase |
|  | 8539288 | C | T | 0.467 | 0.258 | -0.209 | 8.125 | c.1531G>A | p.Val511Ile | AFUN015793 | NA | Carboxylic ester hydrolase |
|  | 8540525 | C | G | 0.311 | 0.273 | -0.039 | 8.127 | c.365G>C | p.Gly122Ala | AFUN015793 | NA | Carboxylic ester hydrolase |
|  | 8541537 | C | T | 0.5 | 0.261 | -0.239 | 8.129 | c.1354G>A | p.Gly452Ser | AFUN008357 | NA | Cytochrome P450 |
|  | 8541623 | A | G | 0.446 | 0.253 | -0.193 | 8.129 | c.1268T>C | p.Ile423Thr | AFUN008357 | NA | Cytochrome P450 |
|  | 8542022 | A | T | 0.725 | 0.547 | -0.178 | 8.13 | c.945T>A | p.Asp315Glu | AFUN008357 | NA | Cytochrome P450 |
|  | 8545954 | T | G | 0.774 | 0.333 | -0.441 | 8.137 | c.196A>C | p.Lys66Gln | AFUN015792 | CYP6P9A | cytochrome P450 |
|  | 8564352 | G | C | 0.769 | 0.416 | -0.354 | 8.183 | c.78C>G | p.His26Gln | AFUN019365 | NA | Cytochrome P450 |
|  | 8565634 | C | G | 0.709 | 0.273 | -0.436 | 8.186 | c.1172G>C | p.Arg391Thr | AFUN015802 | CYP6P1 | Cytochrome P450 |
|  | 8566163 | T | C | 0.726 | 0.378 | -0.348 | 8.188 | c.712A>G | p.Ile238Val | AFUN015802 | CYP6P1 | Cytochrome P450 |
|  | 8567494 | A | C | 0.571 | 0.213 | -0.358 | 8.191 | c.1443T>G | p.Asp481Glu | AFUN015801 | CYP6P2 | Cytochrome P450 |
|  | 8568309 | A | G | 0.71 | 0.308 | -0.402 | 8.193 | c.701T>C | p.Leu234Ser | AFUN015801 | CYP6P2 | Cytochrome P450 |
| 3 prime UTR | 8544509 | G | A | 0.676 | 0.362 | -0.313 | 8.134 | c.*55C>T |  | AFUN015792 | CYP6P9A | cytochrome P450 |
|  | 8554326 | C | T | 0.667 | 0.225 | -0.441 | 8.156 | c.*16G>A |  | AFUN015889 | CYP6P9b | cytochrome P450 |
|  | 8533559 | T | A | 0.776 | 0.348 | -0.428 | 8.115 | c.*370A>T |  | AFUN015785 | CYP6AA2 | Cytochrome P450 |
|  | 8533836 | C | T | 0.680 | 0.386 | -0.294 | 8.115 | c.*93G>A |  | AFUN015785 | CYP6AA2 | Cytochrome P450 |
|  | 8531181 | G | T | 0.625 | 0.164 | -0.461 | 8.110 | c.*393C>A |  | AFUN015786 | CYP6AA1 | Cytochrome P450 |
|  | 8531446 | G | A | 0.585 | 0.295 | -0.290 | 8.111 | c.*128C>T |  | AFUN015786 | CYP6AA1 | Cytochrome P450 |
|  | 8569297 | C | A | 0.632 | 0.354 | -0.278 | 8.196 | c.*250G>T |  | AFUN015714 | CYP6AD1 | Cytochrome P450 |
|  | 8560084 | T | C | 0.746 | 0.267 | -0.480 | 8.171 | c.*356A>G |  | AFUN020895 | NA | Cytochrome P450 |
|  | 8560279 | C | T | 0.717 | 0.321 | -0.396 | 8.172 | c.*161G>A |  | AFUN020895 | NA | Cytochrome P450 |
| stop gained | 8640035 | C | T | 0.212 | 0.224 | 0.012 | 8.382 | c.325C>T | p.Gln109* | AFUN021913 | NA | cathepsin F |
|  | 8640035 | C | T | 0.212 | 0.224 | 0.012 | 8.382 | c.325C>T | p.Gln109* | AFUN014894 | NA | unspecified product |

**Table S11. SNPs in the *rap1* QTL identified by BSA in the F7 mixed cross family are potentially implicated in resistance.** In the F7 mixed cross family, we identified 12 missense SNPs in 8 different genes

| **Effect** | **POS** | **REF** | **ALT** | **SNPindex.LOW** | **SNPindex.HIGH** | **deltaSNP** | **negLog10Pval** | **codon** | **Amino acid** | **ID** | **Name** | **description** |
| --- | --- | --- | --- | --- | --- | --- | --- | --- | --- | --- | --- | --- |
| Missense variant | 8536134 | G | T | 0.729 | 0.212 | -0.516 | Inf | c.1372C>A | p.Leu458Ile | AFUN015787 | NA | Carboxylic ester hydrolase |
|  | 8540480 | A | T | 0.864 | 0.483 | -0.381 | Inf | c.410T>A | p.Phe137Tyr | AFUN015793 | NA | Carboxylic ester hydrolase |
|  | 8532177 | G | A | 0.805 | 0.150 | -0.655 | Inf | c.1013C>T | p.Ala338Val | AFUN015786 | CYP6AA1 | Cytochrome P450 |
|  | 8532483 | C | T | 0.718 | 0.119 | -0.599 | Inf | c.707G>A | p.Arg236Gln | AFUN015786 | CYP6AA1 | Cytochrome P450 (Fragment) |
|  | 8565634 | C | G | 0.849 | 0.220 | -0.630 | Inf | c.1172G>C | p.Arg391Thr | AFUN015802 | CYP6P1 | Cytochrome P450 (Fragment) |
|  | 8566163 | T | C | 0.808 | 0.167 | -0.641 | Inf | c.712A>G | p.Ile238Val | AFUN015802 | CYP6P1 | Cytochrome P450 (Fragment) |
|  | 8567494 | A | C | 0.716 | 0.133 | -0.583 | Inf | c.1443T>G | p.Asp481Glu | AFUN015801 | CYP6P2 | Cytochrome P450 (Fragment) |
|  | 8568309 | A | G | 0.827 | 0.153 | -0.674 | Inf | c.701T>C | p.Leu234Ser | AFUN015801 | CYP6P2 | Cytochrome P450 (Fragment) |
|  | 8564352 | G | C | 0.788 | 0.253 | -0.535 | Inf | c.78C>G | p.His26Gln | AFUN019365 | NA | Cytochrome P450 (Fragment) |
|  | 8542071 | T | C | 0.480 | 0.222 | -0.257 | Inf | c.896A>G | p.Asn299Ser | AFUN008357 | NA | Cytochrome P450 |
|  | 8542176 | C | T | 0.444 | 0.097 | -0.348 | Inf | c.791G>A | p.Arg264His | AFUN008357 | NA | Cytochrome P450 |
|  | 8542719 | A | G | 0.663 | 0.216 | -0.447 | Inf | c.248T>C | p.Met83Thr | AFUN008357 | NA | Cytochrome P450 |
| Stop gained | 8640035 | C | T | 0.125 | 0.478 | 0.353 | Inf | c.325C>T | p.Gln109* | AFUN021913 | NA | cathepsin F |
|  | 8541699 | G | A | 0.475 | 0.179 | -0.296 | Inf | c.1192C>T | p.Gln398* | AFUN008357 | NA | Cytochrome P450 |
|  | 8541699 | G | A | 0.475 | 0.179 | -0.296 | Inf | c.1192C>T | p.Gln398* | AFUN020405 | NA | solute carrier family 8 (sodium/calcium exchanger) |
|  | 8640035 | C | T | 0.125 | 0.478 | 0.353 | Inf | c.325C>T | p.Gln109* | AFUN014894 | NA | unspecified product |
| 3 prime UTR variants | 8544517 | G | C | 0.707 | 0.152 | -0.555 | Inf | c.*47C>G |  | AFUN015792 | CYP6P9A | cytochrome P450 |
|  | 8554336 | T | A | 0.824 | 0.243 | -0.580 | Inf | c.*6A>T |  | AFUN015889 | CYP6P9b | cytochrome P450 |
|  | 8533559 | T | A | 0.867 | 0.305 | -0.563 | Inf | c.*370A>T |  | AFUN015785 | CYP6AA2 | Cytochrome P450 (Fragment) |
|  | 8533836 | C | T | 0.843 | 0.288 | -0.555 | Inf | c.*93G>A |  | AFUN015785 | CYP6AA2 | Cytochrome P450 (Fragment) |
|  | 8531181 | G | T | 0.794 | 0.246 | -0.547 | Inf | c.*393C>A |  | AFUN015786 | CYP6AA1 | Cytochrome P450 (Fragment) |
|  | 8531446 | G | A | 0.778 | 0.175 | -0.603 | Inf | c.*128C>T |  | AFUN015786 | CYP6AA1 | Cytochrome P450 (Fragment) |
|  | 8569297 | C | A | 0.841 | 0.333 | -0.508 | Inf | c.*250G>T |  | AFUN015714 | CYP6AD1 | Cytochrome P450 (Fragment) |
|  | 8560084 | T | C | 0.826 | 0.224 | -0.602 | Inf | c.*356A>G |  | AFUN020895 | NA | Cytochrome P450 (Fragment) |
|  | 8560279 | C | T | 0.859 | 0.241 | -0.618 | Inf | c.*161G>A |  | AFUN020895 | NA | Cytochrome P450 (Fragment) |
|  | 8541470 | A | G | 0.333 | 0.108 | -0.225 | Inf | c.*32T>C |  | AFUN008357 | NA | Cytochrome P450 |

**Table S12. An overlap of missense SNPs identified from the BSA in the F_7_ isofemale and mixed cross families in key detoxification genes in the *rap1* QTL.** All the missense SNPs identified in the *CYP6* cluster located on the negative strand also cause a missense mutation in the solute carrier family 8 (sodium/calcium exchanger), extending 130kb on the leading strand.

| **Effect** | **POS** | **REF** | **ALT** | **Genen** | **codon** | **Amino Acid** | **Iso DE freq** | **Iso AL freq** | **Mix DE freq** | **Mix AL freq** | **Name** | **description** |
| --- | --- | --- | --- | --- | --- | --- | --- | --- | --- | --- | --- | --- |
| Missense | 8532177 | G | A | AFUN015786 | c.1013C>T | p.Ala338Val | 0.68181818 | 0.24193548 | 0.8045977 | 0.15 | CYP6AA1 | Cytochrome P450 |
|  | 8532483 | C | T | AFUN015786 | c.707G>A | p.Arg236Gln | 0.6031746 | 0.28571429 | 0.71830986 | 0.11940299 | CYP6AA1 | Cytochrome P450 |
|  | 8536134 | G | T | AFUN015787 | c.1372C>A | p.Leu458Ile | 0.54545455 | 0.31944444 | 0.72857143 | 0.21212121 | NA | Carboxylic ester hydrolase |
|  | 8539288 | C | T | AFUN015793 | c.1531G>A | p.Val511Ile | 0.46666667 | 0.25757576 | NA | NA | NA | Carboxylic ester hydrolase |
|  | 8540480 | A | T | AFUN015793 | c.410T>A | p.Phe137Tyr | NA | NA | 0.864 | 0.48314607 | NA | Carboxylic ester hydrolase |
|  | 8540525 | C | G | AFUN015793 | c.365G>C | p.Gly122Ala | 0.31132075 | 0.27272727 | NA | NA | NA | Carboxylic ester hydrolase |
|  | 8541537 | C | T | AFUN008357 | c.1354G>A | p.Gly452Ser | 0.5 | 0.26136364 | NA | NA | NA | Cytochrome P450 |
|  | 8541623 | A | G | AFUN008357 | c.1268T>C | p.Ile423Thr | 0.44578313 | 0.25301205 | NA | NA | NA | Cytochrome P450 |
|  | 8542022 | A | T | AFUN008357 | c.945T>A | p.Asp315Glu | 0.72527473 | 0.54716981 | NA | NA | NA | Cytochrome P450 |
|  | 8542071 | T | C | AFUN008357 | c.896A>G | p.Asn299Ser | NA | NA | 0.47959184 | 0.22222222 | NA | Cytochrome P450 |
|  | 8542176 | C | T | AFUN008357 | c.791G>A | p.Arg264His | NA | NA | 0.44444444 | 0.09677419 | NA | Cytochrome P450 |
|  | 8542719 | A | G | AFUN008357 | c.248T>C | p.Met83Thr | NA | NA | 0.6625 | 0.21568627 | NA | Cytochrome P450 |
|  | 8545954 | T | G | AFUN015792 | c.196A>C | p.Lys66Gln | 0.77419355 | 0.33333333 | NA | NA | CYP6P9A | cytochrome P450 |
|  | 8564352 | G | C | AFUN019365 | c.78C>G | p.His26Gln | 0.76923077 | 0.41558442 | 0.7875 | 0.25274725 | NA | Cytochrome P450 |
|  | 8565634 | C | G | AFUN015802 | c.1172G>C | p.Arg391Thr | 0.70886076 | 0.27272727 | 0.84931507 | 0.21978022 | CYP6P1 | Cytochrome P450 |
|  | 8566163 | T | C | AFUN015802 | c.712A>G | p.Ile238Val | 0.72619048 | 0.37804878 | 0.80769231 | 0.16666667 | CYP6P1 | Cytochrome P450 |
|  | 8567494 | A | C | AFUN015801 | c.1443T>G | p.Asp481Glu | 0.57142857 | 0.21333333 | 0.71621622 | 0.13333333 | CYP6P2 | Cytochrome P450 |
|  | 8568309 | A | G | AFUN015801 | c.701T>C | p.Leu234Ser | 0.71014493 | 0.30769231 | 0.82716049 | 0.15277778 | CYP6P2 | Cytochrome P450 |

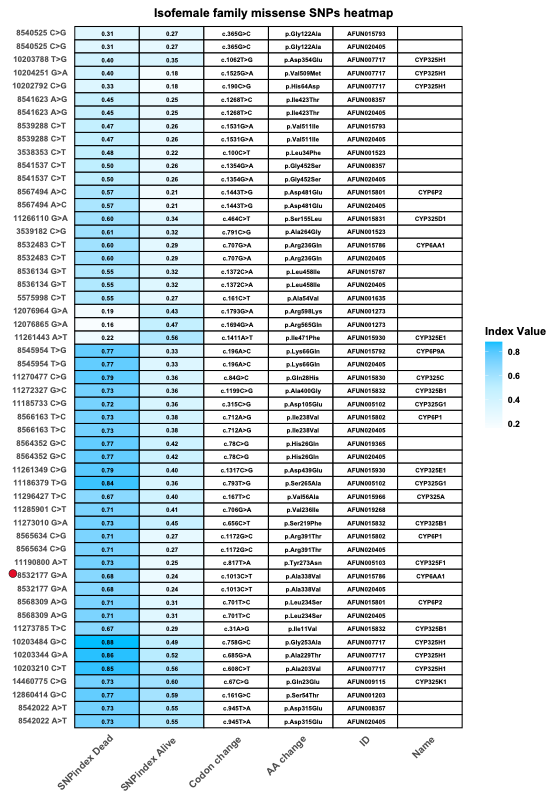

**Figure S10. A heatmap of missense SNPs identified from the BSA analysis in the F_7_ isofemale family. Missense SNPs were identified notably in the *CYP6* and the *CYP32* cluster. The red dot highlights is a missense SNP that was also a marker identified in the peak of the conventional QTL mapping using ddRADseq.**

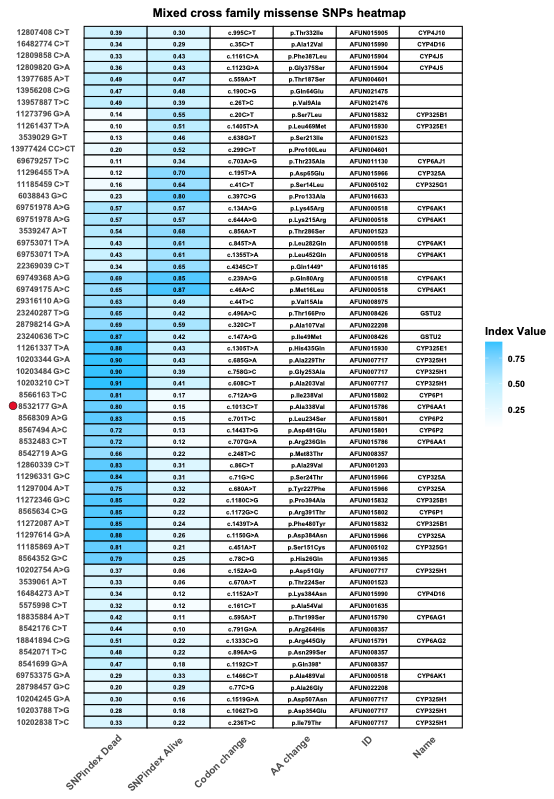

**Figure S11. A heatmap of missense SNPs identified from the BSA analysis in the F_7_ mixed cross family. Missense SNPs identified in the peak of the *rap1* QTL using ddRADseq are highlighted with a red dot.**

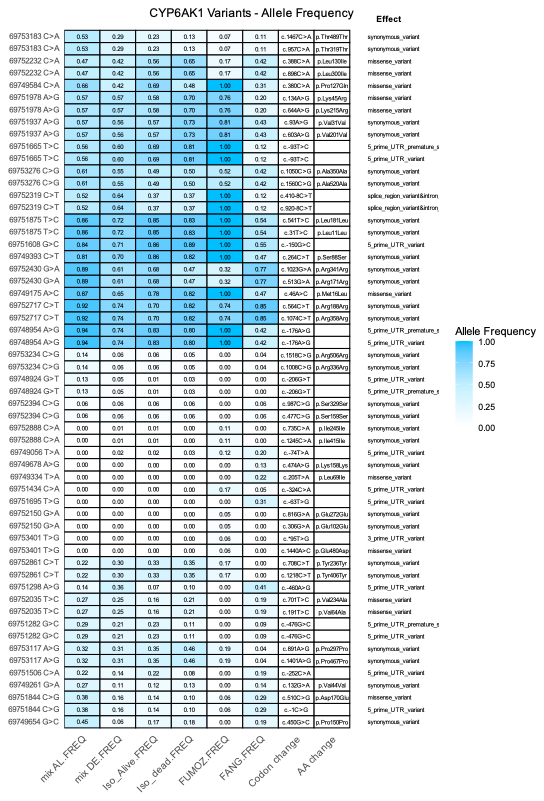

**Figure S12. Allele frequency of SNPs in the *CYP6K1* and their corresponding functional effect.**

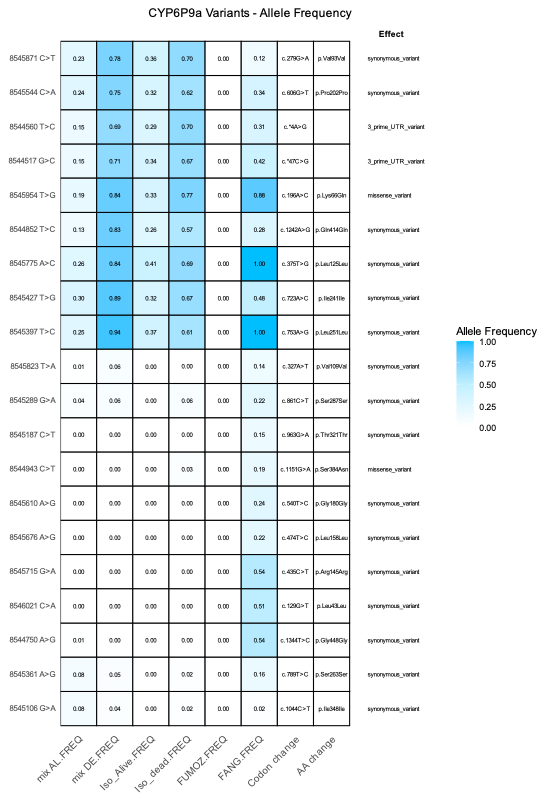

**Figure S13. Allele frequency of SNPs in the *CYP6P9a* and their corresponding functional effect.**

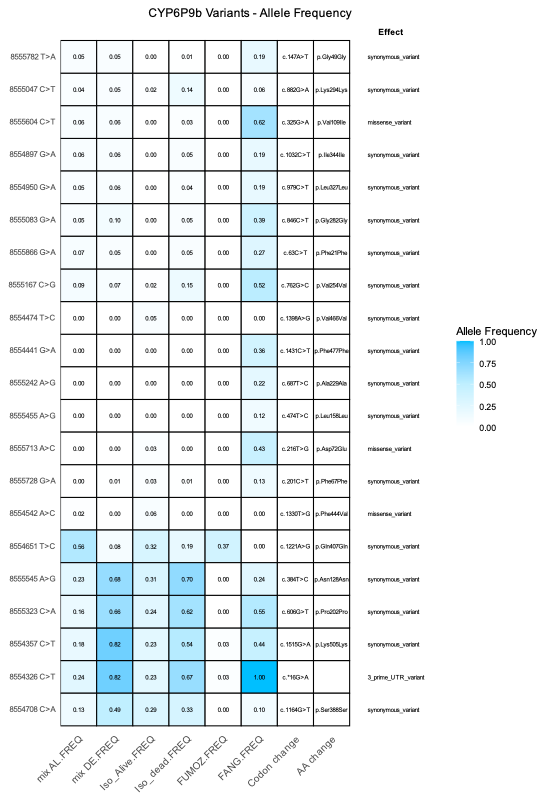

**Figure S14. Allele frequency of SNPs in the *CYP6P9b* and their corresponding functional effect.**

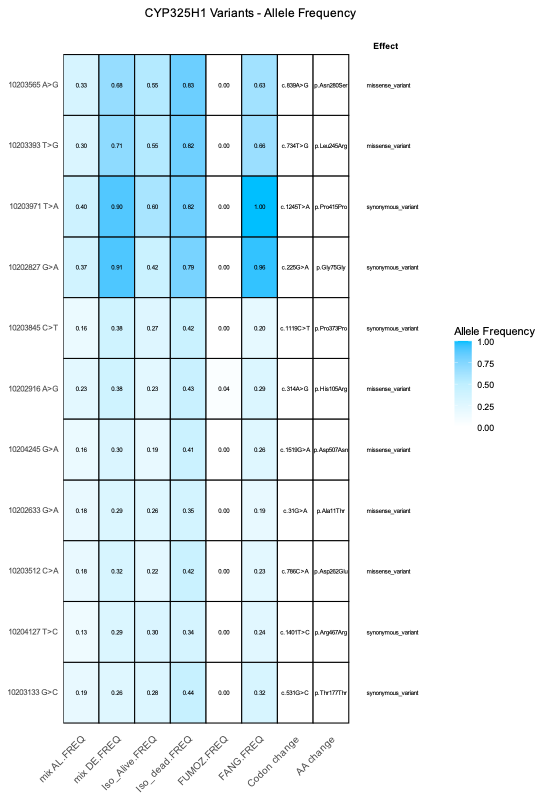

**Figure S15. Allele frequency of SNPs in the *CYP325H1* and their corresponding functional effect.**
